# Myotubularin-related phosphatase 5 is a critical determinant of autophagy in neurons

**DOI:** 10.1101/2021.07.20.453106

**Authors:** Jason P. Chua, Karan Bedi, Michelle T. Paulsen, Mats Ljungman, Elizabeth M. H. Tank, Erin S. Kim, Jennifer M. Colón-Mercado, Michael E. Ward, Lois S. Weisman, Sami J. Barmada

## Abstract

Autophagy is a conserved, multi-step process of capturing proteolytic cargo in autophagosomes for lysosome degradation. The capacity to remove toxic proteins that accumulate in neurodegenerative disorders attests to the disease-modifying potential of the autophagy pathway. However, neurons respond only marginally to conventional methods for inducing autophagy, limiting efforts to develop therapeutic autophagy modulators for neurodegenerative diseases. The determinants underlying poor autophagy induction in neurons and the degree to which neurons and other cell types are differentially sensitive to autophagy stimuli are incompletely defined. Accordingly, we sampled nascent transcript synthesis and stabilities in fibroblasts, induced pluripotent stem cells (iPSCs) and iPSC-derived neurons (iNeurons), thereby uncovering a neuron-specific stability of transcripts encoding myotubularin-related phosphatase 5 (MTMR5). MTMR5 is an autophagy suppressor that acts with its binding partner, MTMR2, to dephosphorylate phosphoinositides critical for autophagy initiation and autophagosome maturation. We found that MTMR5 is necessary and sufficient to suppress autophagy in iNeurons and undifferentiated iPSCs. Using optical pulse labeling to visualize the turnover of endogenously-encoded proteins in live cells, we observed that knockdown of MTMR5 or MTMR2, but not MTMR9, significantly enhances neuronal degradation of TDP-43, an autophagy substrate implicated in several neurodegenerative diseases. Accordingly, our findings establish a regulatory mechanism of autophagy intrinsic to neurons and targetable for clearing disease-related proteins in a cell type-specific manner. In so doing, our results not only unravel novel aspects of neuronal biology and proteostasis, but also elucidate a strategy for modulating neuronal autophagy that could be of high therapeutic potential for multiple neurodegenerative diseases.

## INTRODUCTION

Neurodegenerative diseases belong to a heterogeneous group of sporadic and familial disorders with typical onset in mid- to late-life. These conditions are rapidly rising in prevalence because of increased longevity of the population^1–3^. In spite of diverse clinical manifestations and degeneration in distinct neuroanatomical regions particular for each disease, neurodegenerative disorders harbor overlapping histopathologic features, including the abnormal accumulation of misfolded and aggregated proteins^4^. Although the relationship between such aggregates and the pathogenesis of each disease is incompletely understood, these shared pathologic characteristics implicate age-related or genetic dysfunction of protein quality control mechanisms as convergent pathways to neurodegeneration.

One such mechanism of protein quality control is macroautophagy, here referred to as autophagy. From Greek etymologic roots meaning “self-digestion,” autophagy is a highly conserved, multi-step process of capturing protein and organelle substrates, both selectively and in bulk, into specialized autophagosome vesicles for trafficking to lysosomes, within which such cargo is degraded by proteolytic enzymes. Autophagy operates at a constitutive level but can be stimulated as part of the adaptive response to stress and nutrient deprivation to maintain protein homeostasis. Precise and multifaceted regulatory machinery is required for coordinating autophagy induction. Such regulation is accomplished by (i) signaling cascades inhibiting the autophagy-suppressive mTOR pathway^5,6^, (ii) kinases synthesizing phosphoinositide scaffolds upon which autophagy initiation complexes are assembled^7,8^, and (iii) additional multimeric proteins directing autophagosome membrane elongation and autophagosome-lysosome fusion^9,10^.

Many of the aggregation-prone proteins found in neurodegenerative diseases are autophagy substrates. Furthermore, genetic ablation of autophagy components produces neurodegeneration in mice^11,12^, and inherited forms of neurodegenerative disorders are caused by, or associated with, mutations in key autophagy-related genes, including *SQSTM1* (sequestosome-1 or p62)^13^, *TBK1* (TANK binding kinase 1)^14^, and *OPTN* (optineurin)^15^ in familial amyotrophic lateral sclerosis; *PRKN* (Parkin)^16^ and *PINK1* (PTEN-induced kinase 1)^17,18^ in Parkinson disease; *VCP* (valosin containing protein)^19^ in multisystem proteinopathy; *WDR45* (WD-repeat domain 45, also known as WIPI4)^20^ in β-propeller protein-associated neurodegeneration (BPAN); and *EPG5* (Ectopic p-granules autophagy protein 5 homolog)^21^ in Vici syndrome. Collectively, these observations suggest that intact autophagy is required for maintaining neuronal proteostasis and preventing neurodegeneration. The susceptibility of pathogenic proteins to autophagy and the neuroprotection provided by normal autophagy function, together with growing evidence demonstrating that autophagic degradation of these proteins rescues cellular toxicity^22–38^, implies that the autophagy pathway is a promising therapeutic target for the broad category of neurodegenerative disease. Despite this, however, autophagy modulators have largely failed to provide clinically meaningful benefit for those with neurodegenerative disorders^39–47^. Additionally, the use of currently available modulators of autophagy is limited by narrow therapeutic indices, dose-dependent toxicity, and wide-ranging adverse effects due to pleiotropic target engagement^6,48–51^.

The inherent resistance of neurons to conventional methods of autophagy induction may also contribute to the inefficacy of autophagy stimulators tested in clinical trials. Starvation and mTOR inhibition are potent inducers of autophagy in most cell types but are largely ineffective in neurons, despite successful target engagement^52–54^. The negative results in clinical trials thus far may therefore relate to inadequate induction of neuronal autophagy^52,54–56^, and alternative strategies for augmenting autophagy in neurons that overcome the prevailing barriers against autophagy-based therapies are sorely needed. Nevertheless, the critical mechanisms that underlie the insensitivity of neurons to autophagy stimuli and would be amenable for therapeutic targeting have yet to be identified.

Here, we sought to uncover neuronal determinants governing autophagy and leading to the relative resistance of neurons to autophagy inducers. Using unbiased, genome-wide assessments of cell type-specific gene expression, we identified a selective enrichment of myotubularin-related phosphatase 5 (MTMR5) in iNeurons, or neurons derived from human induced pluripotent stem cells (iPSCs). MTMR5, also known as SET binding factor 1 (SBF1), belongs to a 14-member family of myotubularin-related phosphatases (MTMRs) that catalyze the removal of phosphate groups from the third and fifth positions on membrane phosphoinositides, including phosphatidylinositol-3-phosphate and phosphatidylinositol-3,5-bisphosphate (PtdIns3P and PtdIns(3,5)P2)^57,58^. Since these enzymatic events prevent recruitment of autophagy initiation complexes and reduce autophagy induction, MTMRs are considered autophagy suppressors^7,59,60^. Notably, MTMR5 is catalytically inactive^59,60^, and instead associates with its paralog and active phosphatase, MTMR2, as a heterodimer to regulate MTMR2’s localization and enhance its phosphatase activity^61^. Similar to MTMR2^62^ and MTMR13^63^, MTMR5 is essential for maintaining peripheral nerves, and loss-of-function mutations lead to dysmyelination and a subtype of Charco-Marie-Tooth disease, CMT4B^64,65^. However, the precise role of MTMR5 in the central nervous system, and whether MTMR5 regulates autophagy to similar extents in different cell types, is unknown. In this study, our identification of MTMR5’s selective enrichment in neurons hinted at a determinative role of MTMR5 in regulating neuronal autophagy. We therefore sought to manipulate MTMR5 expression and found that reductions in MTMR5 robustly enhanced neuronal autophagy, while neuron-like insensitivity to autophagy induction was recapitulated by MTMR5 overexpression in non-neuronal cells. In so doing, our results unravel not only a pivotal function of myotubularins in neuronal proteostasis, but also establish a novel target for potentiating autophagy in neurons for therapy design in neurodegenerative diseases.

## RESULTS

### Neurons are resistant to autophagy induction by Torin1

We first assessed whether neurons generate autophagosomes to a similar degree as non-neuronal cell types. To monitor autophagosome biogenesis non-invasively, we used an iPSC line edited with CRISPR/Cas9 to tag endogenous LC3 — a macroautophagy substrate as well as a marker of developing autophagosomes — with mEGFP at the N-terminus (Fig. 1A). We subsequently edited this iPSC line further to allow doxycycline-inducible, rapid differentiation of nearly 100% efficiency^66^ into glutamatergic forebrain-like neurons using TALENS (iNeurons; Fig. 1B-C), or a piggybac/transposase system for doxycycline-inducible differentiation into skeletal muscle^67^ (iMuscle; Fig. 1D, S1). Lastly, we differentiated mEGFP-LC3 iPSCs into astrocytes using dual-SMAD inhibition, as previously described^68^ (iAstrocytes; Fig. 1C). Treatment with 250nM Torin1 for 4h produced large numbers of mEGFP-positive autophagosomes in iAstrocytes, iMuscle, and undifferentiated iPSCs, but significantly fewer autophagosomes in iNeurons (Fig. 1E-G). These results demonstrate that, compared to non-neuronal cells, neurons are less sensitive to the autophagy-inducing effects of the mTOR inhibitor Torin1.

**Figure 1.**
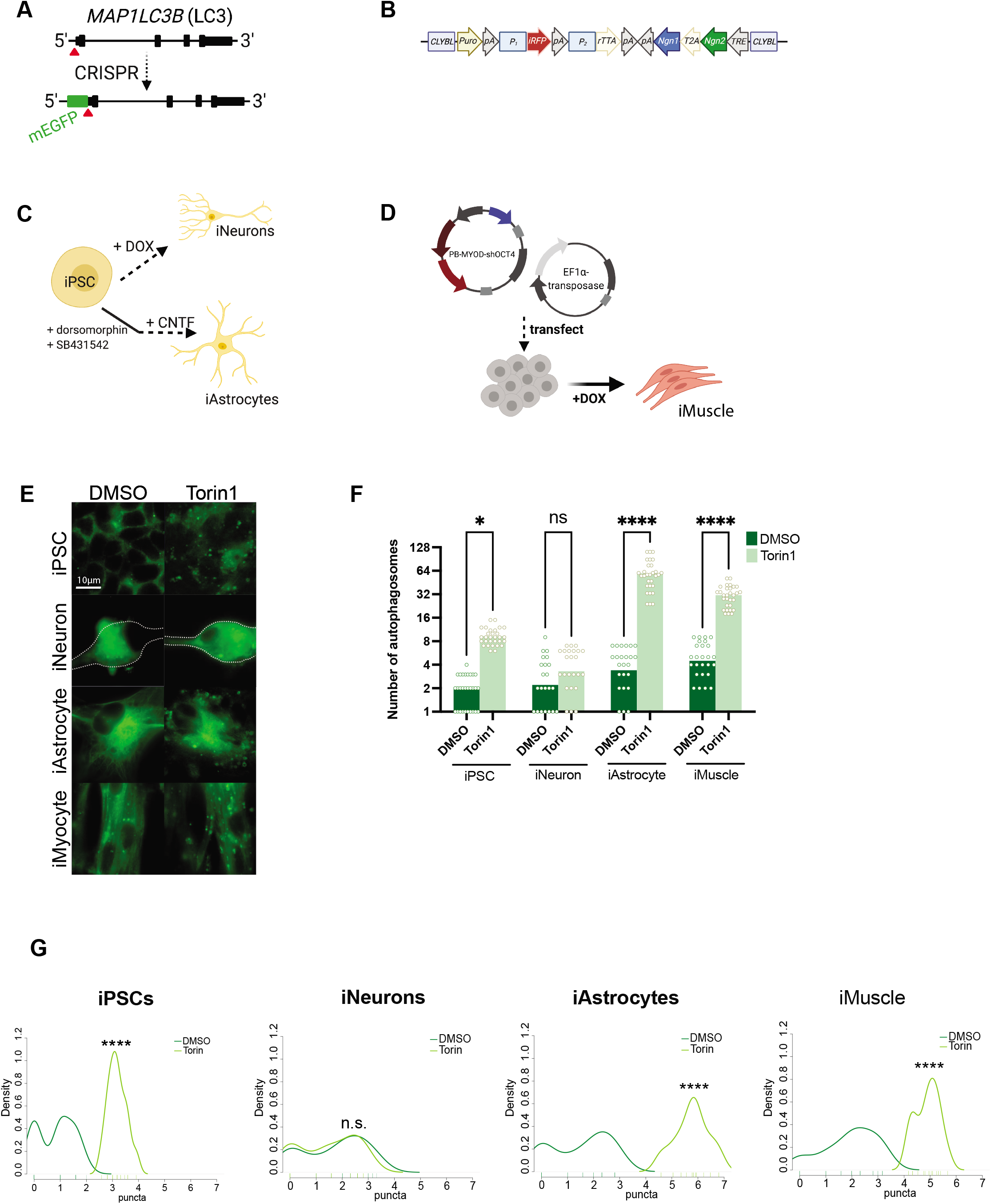
Neurons are resistant to Torin1-mediated induction of autophagy. (**A**) Targeting strategy using CRISPR/Cas9 to knock-in mEGFP immediately 5’ to exon1 in the *MAP1LC3B* gene. (**B**) Schematic of the cassette used to integrate *NGN1* and *NGN2* at the *CLYBL* safe harbor locus under the control of a Tet-ON system. *Puro*, puromycin-resistance gene; *pA*, poly-A tail; *P_1_, P_2_,* promotors; *iRFP*, near-infrared fluorescent protein; *rTTA*, reverse tetracycline-controlled transactivator; *NGN1* and *NGN2*, neurogenin-1 and −2; *T2A*, self-cleaving peptide; *TRE*, tetracycline response element. (**C**) Protocols used to differentiate iPSCs into iNeurons using doxycycline-mediated, forced expression of differentiation factors, or into iAstrocytes using dual-SMAD inhibition followed by terminal differentiation by culturing in CNTF. (**D**) Protocol for differentiating iPSCs into iMuscle using a piggybac/transposase system to integrate doxycycline-inducible *MYOD* and *OCT4* shRNA. (**E**) Representative images of mEGFP-LC3-positive vesicles in the cell types indicated after treatment with DMSO vehicle or 250nM Torin1 for 4h. (**F**) Scatterplots of blinded manual quantifications of mEGFP-LC3-positive vesicles imaged as in (E). Data are from three independent experiments. n.s., not significant; *p<0.05; ****p<0.0001; one-way ANOVA with Šídák’s multiple comparisons test. (**G**) Density plots of data from (F). n.s., not significant; ****p<0.0001, two-sample Kolmogorov-Smirnov test.

### MTMR5 expression is selectively enhanced in neurons

To uncover differences across the human genome in the expression of autophagy-related genes that may account for the relative insensitivity of neurons to mTOR inhibition, we analyzed the transcriptome of iNeurons, and compared this to isogenic iPSCs and the fibroblasts from which these iPSCs were reprogrammed. Steady-state measurements of mRNA transcripts via RNA-seq or PCR-based methods do not capture key aspects of gene expression such as rates of synthesis and turnover of RNA^69,70^. Therefore, we took advantage of Bru-seq and BruChase-seq to assess the relative synthesis and stabilities of RNA transcripts genome-wide. Briefly, these methods consist of the incorporation of bromouridine (BrU) into synthesizing RNA, followed with or without a chase using excess uridine. BrU-labeled transcripts are pulled down with anti-bromouridine antibodies, synthesized into cDNA, then analyzed by deep sequencing (Fig. 2A)^69,70^.

**Figure 2.**
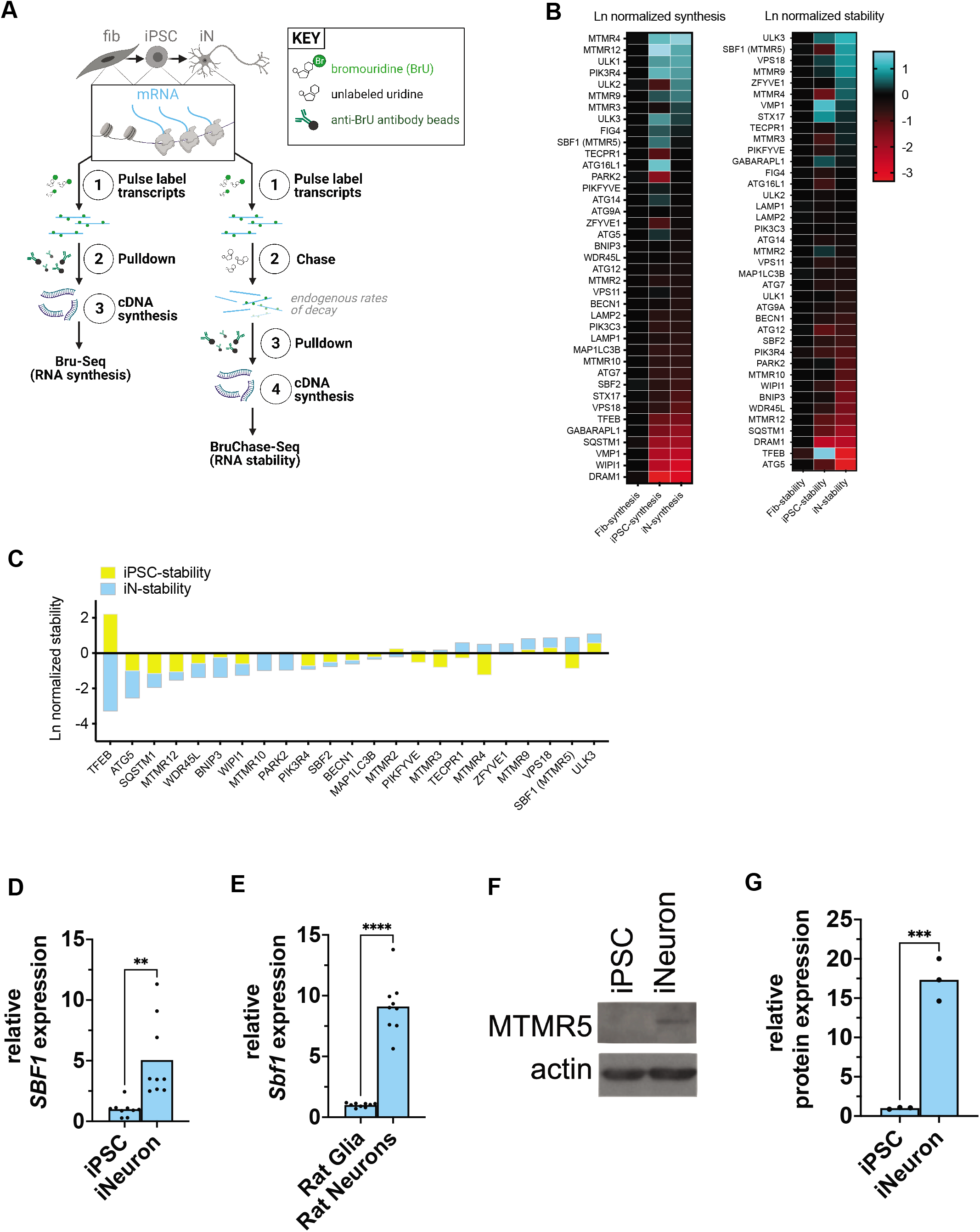
MTMR5 is enriched in neurons. (**A**) Schematic of Bru-Seq and Bru-Chase-Seq. RNA transcripts from fibroblasts, iPSCs reprogrammed from the fibroblasts, and iNeurons differentiated from the same iPSCs were pulse-labeled with bromouridine with and without chasing with unlabeled uridine, followed by pulldown with anti-bromouridine antibody beads, then subjected to RNA-Seq. (**B**) Heatmaps of RNA synthesis data from Bru-Seq (left) and RNA stability data from BruChase-Seq (right), plotted for each cell type and natural log (Ln)-normalized to fibroblasts. Fib, fibroblast; iPSC, induced pluripotent stem cell; iN, iNeuron. (**C**) Bar graph of RNA stability data from BruChase-Seq, Ln-normalized to fibroblasts. *SBF1* was among the most stable RNA transcripts. (**D**) RT-PCR measurements of total steady-state *SBF1* RNA in iNeurons compared to iPSCs. **p<0.01, Student’s *t* test. (**E**) RT-PCR measurements of total steady-state *Sbf1* RNA in primary rat neurons compared to rat glia. ****p<0.0001, Student’s *t* test. (**F**) Representative Western analysis of MTMR5 protein and actin loading control in iNeurons compared to iPSCs. (**G**) Band intensity quantifications of MTMR5 as depicted in (F), normalized to actin band intensity. Data are from three independent experiments. ***p<0.001, Student’s *t* test.

Among several candidate transcripts, *SBF1*—encoding the protein MTMR5^61,64,71^—exhibited significantly higher stability in iNeurons compared to fibroblasts and undifferentiated iPSCs (Fig. 2B-C). *SBF1* transcription rates were similar across cell types, indicating a selective difference in mRNA stability for *SBF1*. In contrast, the stability of *MTMR2* was not significantly different in the three cell types (Fig. 2B-C). We then asked if the heightened stability of *SBF1* correlated with total steady state levels of *SBF1* RNA. Consistent with our Bru-seq data, measurements of *SBF1* RNA by RT-PCR revealed a selective, nearly five-fold enrichment in iNeurons compared to their iPSCs of origin (Fig. 2D). This expression pattern was conserved across species, as rat cortical neurons exhibited a similar, higher abundance of orthologous *Sbf1* RNA compared to cortical glia (Fig. 2E). These results correlated well with multiple and independent RNA-seq datasets showing higher expression of *SBF1* in neurons (Fig. S2A). When comparing against other, additional cell types, expression of neuronal *SBF1* was also higher compared to iMuscle *SBF1*, though similarly enriched in iAstrocytes (Fig. S2B). For *MTMR2*, steady-state RNA levels were only slightly elevated in iNeurons and rat cortical neurons, but approximately 50% lower in iAstrocytes and iMuscle compared to undifferentiated iPSCs (Fig. S2C). Importantly, neuronal enrichment for *SBF1* was not limited to RNA alone—western blots also demonstrated higher levels of MTMR5 protein in iNeurons compared to undifferentiated iPSCs (Fig. 2F-G). Together, these data corroborate our Bru-seq and BruChase-seq analyses and validate a neuron-specific excess of the autophagy suppressor, MTMR5, due to enhanced stability of the *SBF1* transcript.

### MTMR5 is sufficient for suppressing autophagy

We next wondered if enhancing MTMR5 expression leads to a neuron-like blunting of autophagy induction in non-neuronal cells. We chose to increase endogenous MTMR5 expression through CRISPR activation (CRISPR_A_), in which catalytically inactive Cas9 (dCas9) fused to transcriptional activators (VP64, MS2, p65, HSF1) is directed to a locus of interest by a single guide RNA (sgRNA) to enhance native gene expression (Fig. 3A)^72,73^. Using the CRISPR_A_ system and a sgRNA specific for the *SBF1* locus, we increased expression of endogenous MTMR5 in undifferentiated iPSCs (Fig. 3B). We then measured the effect of MTMR5 overexpression on autophagosome number after Torin1-mediated induction of autophagy. iPSCs expressing non-targeted sgRNA showed no change in autophagosome biogenesis after treatment with Torin1 (Fig. 3C-E). However, cells expressing sgRNA targeting *SBF1* (Fig. 3C-E) demonstrated a significant attenuation of autophagosome induction after Torin1 treatment, similar to that observed in iNeurons (Fig. 1E-G). These results confirm that MTMR5 is sufficient for restricting autophagosome biogenesis, as its overexpression in non-neuronal cells recapitulates the insensitivity of neurons to Torin1-mediated induction of autophagy.

**Figure 3.**
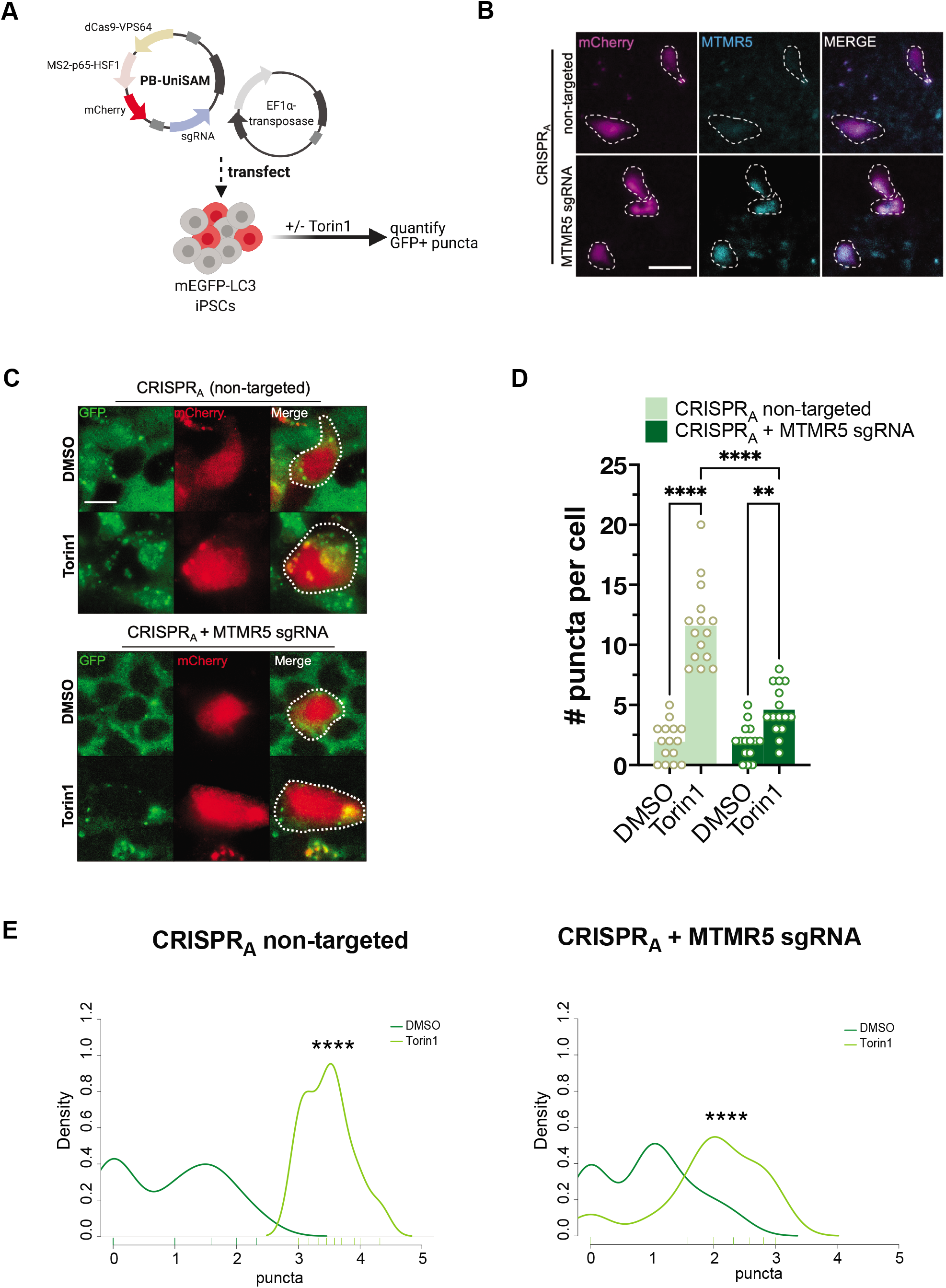
MTMR5 is sufficient to desensitize iPSCs to Torin1 induction of autophagy. (**A**) Schematic of CRISPR_A_ experimental workflow using piggybac/transposase system to express dCas9 fused with VPS64-MS2-p65 transactivators. Cells were treated with 250nM Torin1 for 4h then fixed, stained, and imaged. (**B**) Representative images of iPSCs transfected with CRISPR_A_ vectors without a targeting sgRNA (top) or with sgRNA targeting the native *SBF1* locus, followed by immunocytochemistry staining for MTMR5 to confirm protein overexpression. Cells harboring the piggybac/transposase vectors are indicated by the mCherry expression marker. Scale bar, 15μm. (**C**) Representative images of mEGFP-LC3-positive vesicles visualized in mCherry-positive cells after treatment with DMSO vehicle or Torin1. Scale bar, 10μm. (**D**) Scatterplot of blinded manual quantifications of mEGFP-LC3-positive vesicles imaged as in (C). Data are from three independent experiments. **p<0.01; ****p<0.0001, one-way ANOVA with Šídák’s multiple comparison’s test. (**E**) Density plots of data from (D). ****p<0.0001, two-sample Kolmogorov-Smirnov test.

### MTMR5 is necessary for neuronal resistance to autophagy stimuli

We next asked whether reducing MTMR5 levels augments autophagy induction in neurons. To address this, we transduced mEGFP-LC3 iNeurons with shRNA directed against *SBF1* or a non-targeted control, and visualized mEGFP-LC3 puncta after application of Torin1 (Fig. 4A). We did not observe any changes to autophagosome formation in iNeurons transduced with non-targeted shRNA (Fig. 4B-D). However, knockdown of *SBF1* in iNeurons significantly increased the number of mEGFP-LC3-positive puncta after Torin1 treatment (Fig. 4B-D). As expected, based on the functional interaction of MTMR2 with MTMR5, *MTMR2* knockdown also led to enhanced numbers of puncta after Torin1 treatment (Fig. S3). In contrast, *MTMR9* knockdown failed to produce a similar induction of mEGFP-LC3 puncta in response to Torin1 (Fig. 4B-D). These results indicate that MTMR5 is necessary for repression of autophagy induction in neurons, and the sensitization of neurons to autophagy stimuli is specific to MTMR5.

**Figure 4.**
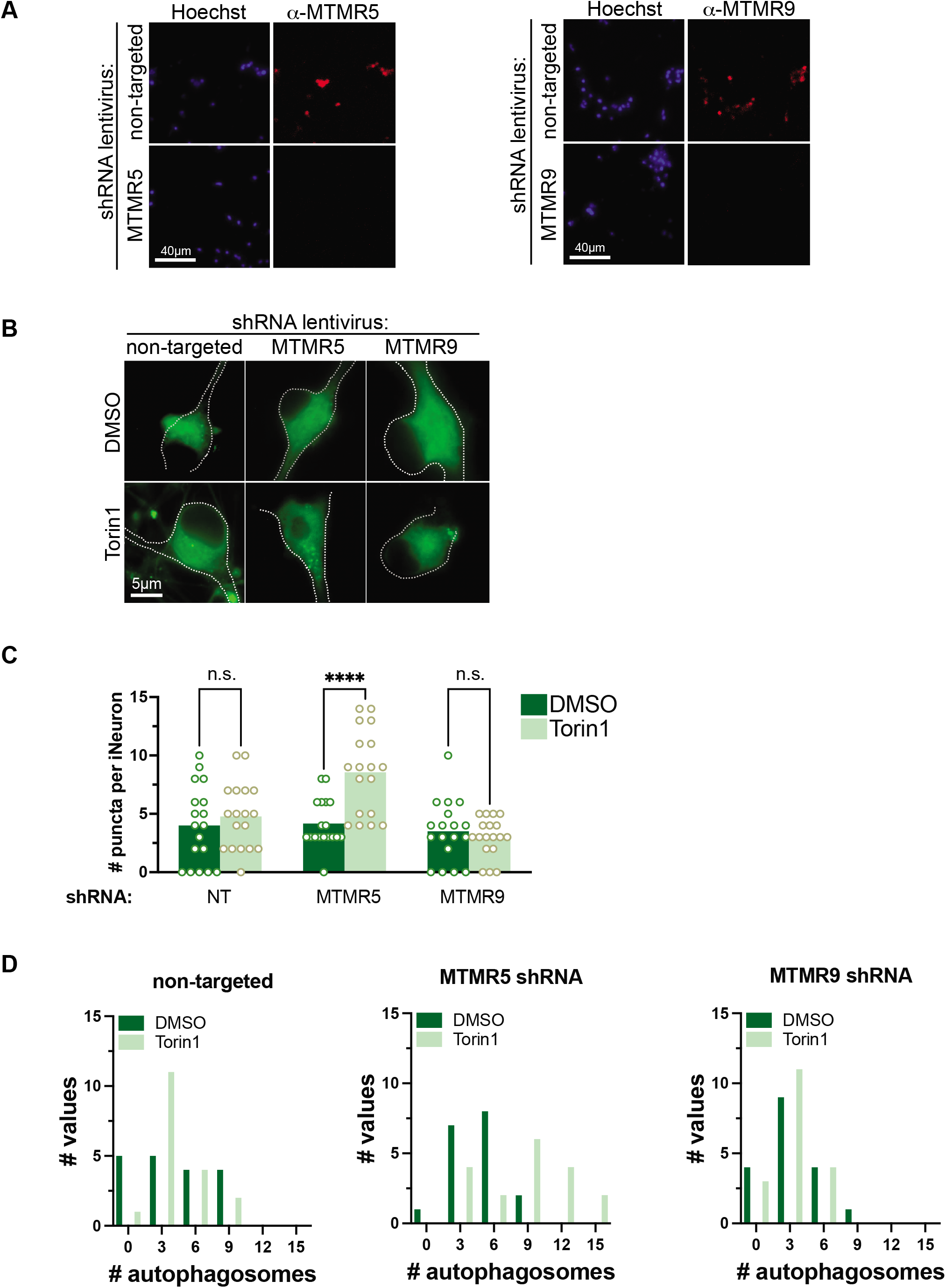
MTMR5 is necessary for suppressing autophagy in neurons. (**A**) Representative immunocytochemical staining against MTMR5 (left) and MTMR9 (right) in iNeurons transduced with non-targeted shRNA lentivirus (top rows), *SBF1* shRNA (bottom left row), or *MTMR9* shRNA lentivirus (bottom right row), respectively. (**B**) Representative images of iNeurons transduced with non-targeted shRNA, *SBF1* shRNA, or *MTMR9* shRNA and treated with DMSO vehicle or 250nM Torin1 for 4h. (**C**) Scatterplots of blinded manual quantifications of mEGFP-LC3-positive puncta imaged in iNeurons as treated in (B). Data are from 3 independent experiments. n.s., not significant; ****p<0.0001, one-way ANOVA with Šídák’s multiple comparisons test. (**D**) Histogram plots of mEGFP-LC3-positive puncta quantifications from (C).

### MTMR5 knockdown enhances degradation through autophagy

Since the accumulation of mEGFP-LC3-positive autophagosomes could indicate autophagy induction or a late-stage block in the pathway, we next asked whether increases in LC3-EGFP number upon *SBF1* knockdown are associated with a corresponding enhancement in autophagic degradation. To address this, we used CRIPSR/Cas9 to label the endogenous protein TDP-43 at the carboxy-terminus with the photoswitchable fluorophore, Dendra2^74,75^ in iPSCs (Fig. 5A). TDP-43 is not only a degradative substrate of autophagy^76,77^, but also integrally involved in the pathogenesis of amyotrophic lateral sclerosis, frontotemporal dementia, and other neurodegenerative proteinopathies^78–83^. Fusion with the Dendra2 allows for non-invasive tracking of endogenous TDP-43 protein levels in live cells via optical pulse labeling (OPL), a technique we previously used to track the turnover of overexpressed TDP-43 in primary neurons^74^. Briefly and as previously described^74^, in OPL we exploit the property of Dendra2 to irreversibly switch emission maxima from green to red wavelengths upon exposure to short-wavelength light^84,85^ (termed “photoconversion,” Fig. 5B). After pulsing the total pool of TDP-43-Dendra2 at the beginning of an experiment, we then use automated fluorescence microscopy^74,86^ to track the pool of photoconverted “red” Dendra2 signal over time (Fig. 5B).

**Figure 5.**
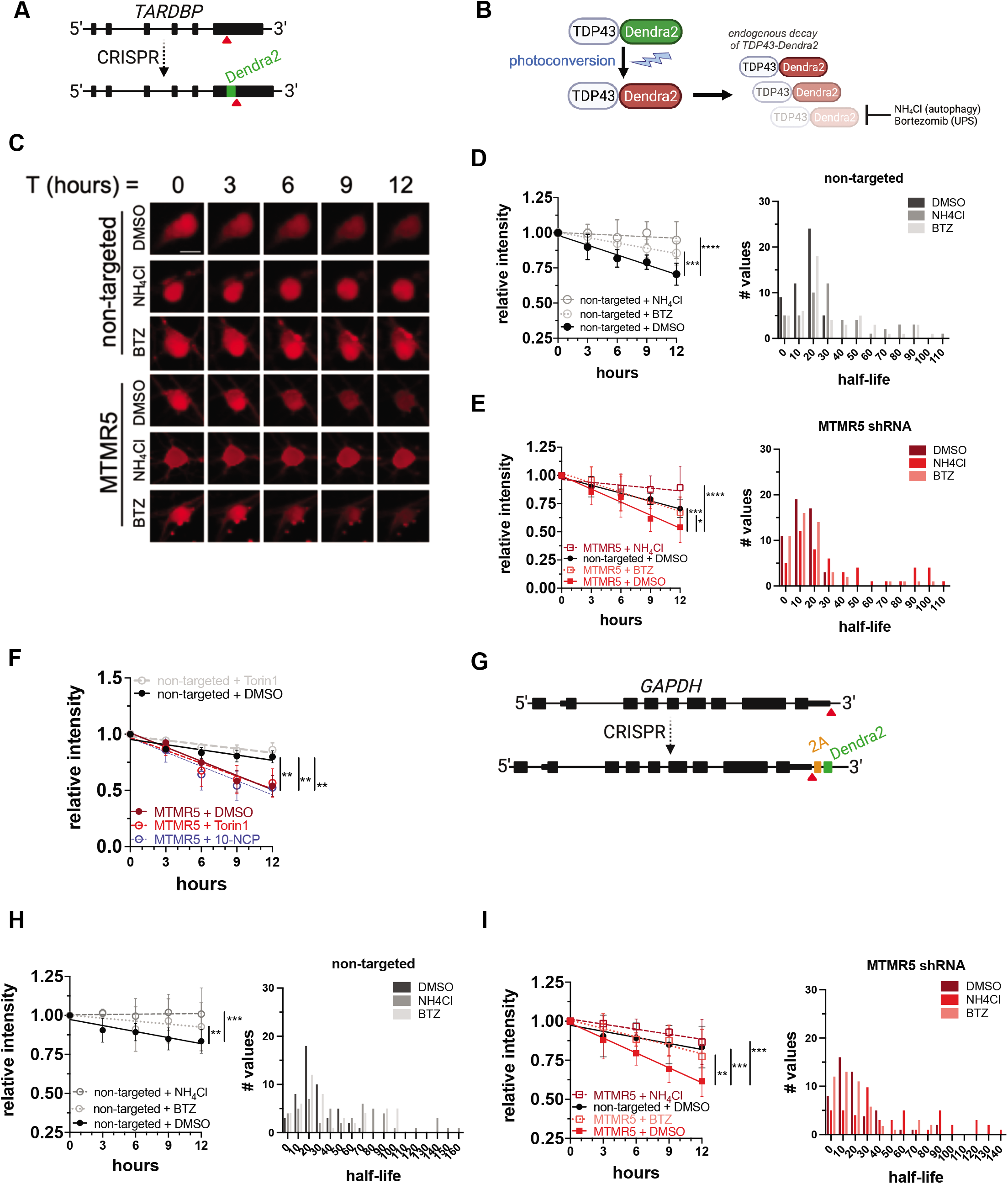
Knockdown of MTMR5 enhances the proteolytic clearance of autophagy substrates. (**A**) Strategy to label endogenous TDP-43 by inserting Dendra2 immediately upstream of the *TARDBP* stop codon using CRISPR/Cas9. (**B**) Schematic of optical pulse labeling (OPL) to photoconvert Dendra2 emission maxima from green to red, followed by tracking red fluorescence decay to monitor TDP-43-Dendra2 degradation, which can be blocked with NH_4_Cl to inhibit autophagy or bortezomib to inhibit the ubiquitin-proteasome system (UPS). (**C**) Representative OPL of TDP-43-Dendra2 iNeurons imaged by automated fluorescence microscopy and after treatment with DMSO, NH_4_Cl, or bortezomib and transduction with non-targeted (top 3 rows) or *SBF1* shRNA lentivirus (bottom 3 rows). Scale bar, 10μm. (**D**-**E**) TDP-43-Dendra2 fluorescence measured in iNeurons transduced with non-targeted (D) or *SBF1* shRNA lentivirus (E) and after the indicated drug treatments (left panels), and histogram plot of the half-lives of each measured iNeuron (right panels). Data are represented at each time point as mean ± SD; *p<0.05; ***p<0.001; ***p<0.001; ****p<0.0001, one-way ANCOVA. (**F**) TDP-43-Dendra2 fluorescence measured in iNeurons transduced with non-targeted or *SBF1* shRNA and after the treatment with DMSO vehicle or autophagy inducing agents Torin1 or 10-NCP. Data are represented at each time point as mean ± SD; **p<0.01, one-way ANCOVA. (**G**) Strategy for inserting Dendra2 at the native *GAPDH* locus using CRISPR/Cas9. 2A, self-cleaving peptide. (**H**-**I**) Dendra2 fluorescence measured in GAPDH-2A-Dendra2 iNeurons transduced with non-targeted (G) or *SBF1* shRNA lentivirus (H) and after the indicated drug treatments (left panels), and histogram plot of the half-lives of each measured iNeuron (right panels). Data are represented at each time point as mean ± SD; **p<0.01; ***p<0.001, one-way ANCOVA.

After transduction with non-targeted shRNA lentivirus, we measured a baseline TDP-43-Dendra2 half-life of approximately 16h in iNeurons via OPL (Fig. 5C, D). TDP-43-Dendra2 half-life was significantly attenuated by inhibiting autophagic flux through treatment with ammonium chloride, and to a lesser extent after proteasomal inhibition with bortezomib (Fig. 5C, D). These observations suggest that TDP-43 undergoes constitutive proteolytic breakdown in iNeurons, primarily through autophagy but also to some degree through the ubiquitin-proteasome pathway. *SBF1* knockdown significantly accelerated TDP-43-Dendra2 decay in iNeurons, resulting in a half-life of approximately 12h (Fig. 5C, E). In the setting of *SBF1* knockdown, treatment with either Torin1 or a neuron-specific inducer of autophagy, 10-NCP^74^, had no further effect on TDP-43-Dendra2 half-life (Fig. 5F), suggesting that *SBF1* knockdown achieved a maximal degree of autophagy stimulation. Consistent with autophagy-mediated degradation of TDP-43-Dendra2 upon *SBF1* knockdown, the protein was effectively stabilized by ammonium chloride treatment (Fig. 5C, E).

We also wondered if the observed changes in protein turnover ascribed to MTMR2/5 knockdown are specific to TDP-43, or if these manipulations affect the autophagic degradation of other proteins. To resolve this, we created a separate iPSC line in which sequences encoding Dendra2 and a self-cleavable 2A peptide were inserted immediately 5’ to the *GAPDH* stop codon (Fig. 5G). These cells, which we termed GAPDH-2A-Dendra2 iPSCs, express untagged Dendra2 that is diffusely distributed throughout the cell, as expected. As measured by OPL, the half-life of Dendra2 in iNeurons was approximately 25h (Fig. 5H). As with TDP-43-Dendra2, treatment with ammonium chloride and bortezomib prolonged the half-life of Dendra2 (Fig. 5H), implying combined autophagic and proteasomal clearance of Dendra2. MTMR5 (Fig. 5I) knockdown accelerated Dendra2 turnover, reducing the half-life to approximately 13h (Fig. 5I). The enhanced decay of Dendra2 was largely abolished with lysosomal inhibition, and less so with proteasomal inhibition (Fig. 5I).

Notably, MTMR5 is a phosphatase-dead member of the myotubularin family^71^, but binds as a heterodimer with its paralog MTMR2, an active phosphatase, to regulate MTMR2’s localization and enhance its activity^61^. We therefore wondered whether MTMR2 serves as a functional effector of MTMR5’s suppressive effect, and if *MTMR2* and *SBF1* knockdown have similar effects on TDP-43-Dendra2 degradation. We performed OPL after transduction with *MTMR2* shRNA and found that, indeed, *MTMR2* knockdown also augmented the degradation of TDP-43-Dendra2 (Fig. 6A, B), resulting in a half-life of about 10h. As before, this effect was blocked by ammonium chloride more so than bortezomib (Fig. 6A, B). In contrast, and in line with our autophagosome quantification experiments (Fig. 4B-D), *MTMR9* knockdown did not significantly change TDP-43-Dendra2 turnover (Fig. 6A, B), indicating that MTMR9 is dispensable for regulating the degradation of autophagy substrates in neurons. Similar to the effects seen with TDP-43-Dendra2, clearance of Dendra2 in GAPDH-2A-Dendra2 iNeurons was also enhanced by knockdown of *MTMR2*, but not *MTMR9* (Fig. 6 C, D). Together, these findings confirm that MTMR5 and MTMR2, but not MTMR9, are primary determinants of autophagy inhibition in iNeurons, and knockdown of either myotubularin increases neuronal degradation of autophagy substrates.

**Figure 6.**
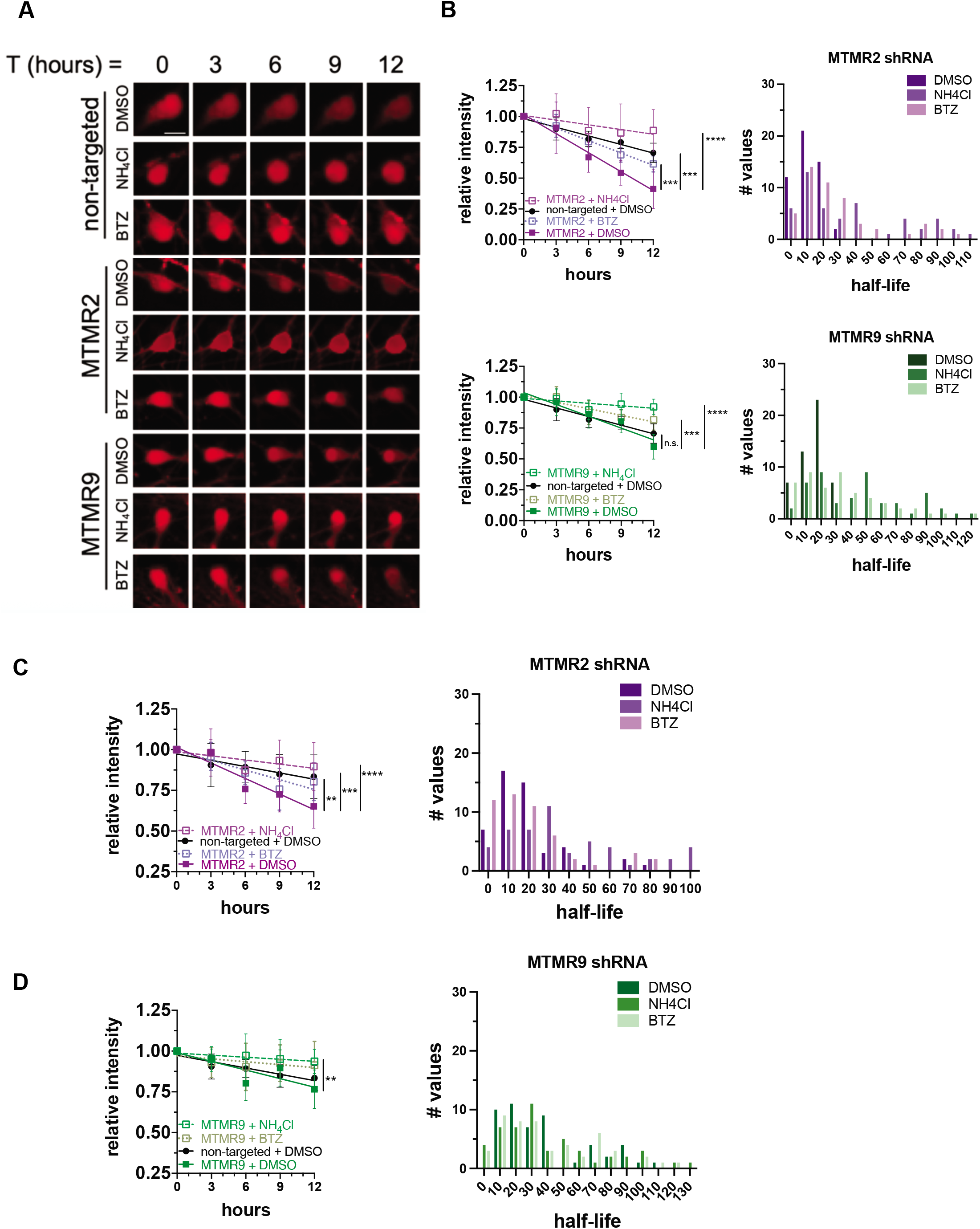
MTMR2, but not MTMR9, is necessary for suppressing neuronal degradation of autophagy substrates. (**A**) Representative OPL of TDP-43-Dendra2 iNeurons imaged by automated fluorescence microscopy and after treatment with DMSO, NH_4_Cl, or bortezomib and transduction with non-targeted (top 3 rows) or *MTMR2* (middle 3 rows), or *MTMR9* (bottom 3 rows) shRNA lentivirus. Scale bar, 10μm. (**B**) TDP-43-Dendra2 fluorescence measured in iNeurons transduced with *MTMR2* (top) or *MTMR9* shRNA lentivirus (bottom) and after the indicated drug treatments (left panels), and histogram plot of the half-lives of each measured iNeuron (right panels). Data are represented at each time point as mean ± SD; n.s., not significant; ***p<0.001; ****p<0.0001, one-way ANCOVA. (**C**-**D**) Dendra2 fluorescence measured in GAPDH-2A-Dendra2 iNeurons transduced with *MTMR2* (C) or *MTMR9* shRNA lentivirus (D) and after the indicated drug treatments (left panels), and histogram plot of the half-lives of each measured iNeuron (right panels). Data are represented at each time point as mean ± SD; **p<0.01; ***p<0.001; ****p<0.0001, one-way ANCOVA.

Although MTMR5 knockdown was sufficient to accelerate the degradation of TDP-43-Dendra2 in glutamatergic iNeurons, this may not be the case for other neuronal subtypes. In addition to forebrain-like glutamatergic neurons, we differentiated iPSCs into motor neurons (iMotor Neurons) through doxycycline-induced expression of *NGN2*, *LHX3*, and *ISL1*, which were integrated at the *CLYBL* safe harbor locus^66,87^ (Fig. S4A). These cells are cultured in different media and exhibit markers of cholinergic lower motor neurons (Fig. S4B). We then asked whether manipulating MTMR5 in iMotor Neurons similarly affected TDP-43-Dendra2 levels by transducing these cells with shRNA lentivirus targeting *SBF1* and measuring TDP-43-Dendra2 turnover by OPL. In cells transduced with non-targeted control, TDP-43-Dendra2 half-life was approximately 12h (Fig. 7A), suggesting a baseline difference in TDP-43-Dendra2 stability between iNeurons and iMotor Neurons. In contrast to what we observed in iNeurons, knockdown of *SBF1*, *MTMR2*, or *MTMR9* in iMotor Neurons had little effect on TDP-43-Dendra2 decay rates (Fig. 7A, C-D upper panels). Together with the relatively rapid turnover of TDP-43-Dendra2 in iMotor Neurons, these results suggest autophagic clearance of TDP-43-Dendra2 is not regulated by MTMR5 in iMotor Neurons, as occurs in iNeurons.

**Figure 7.**
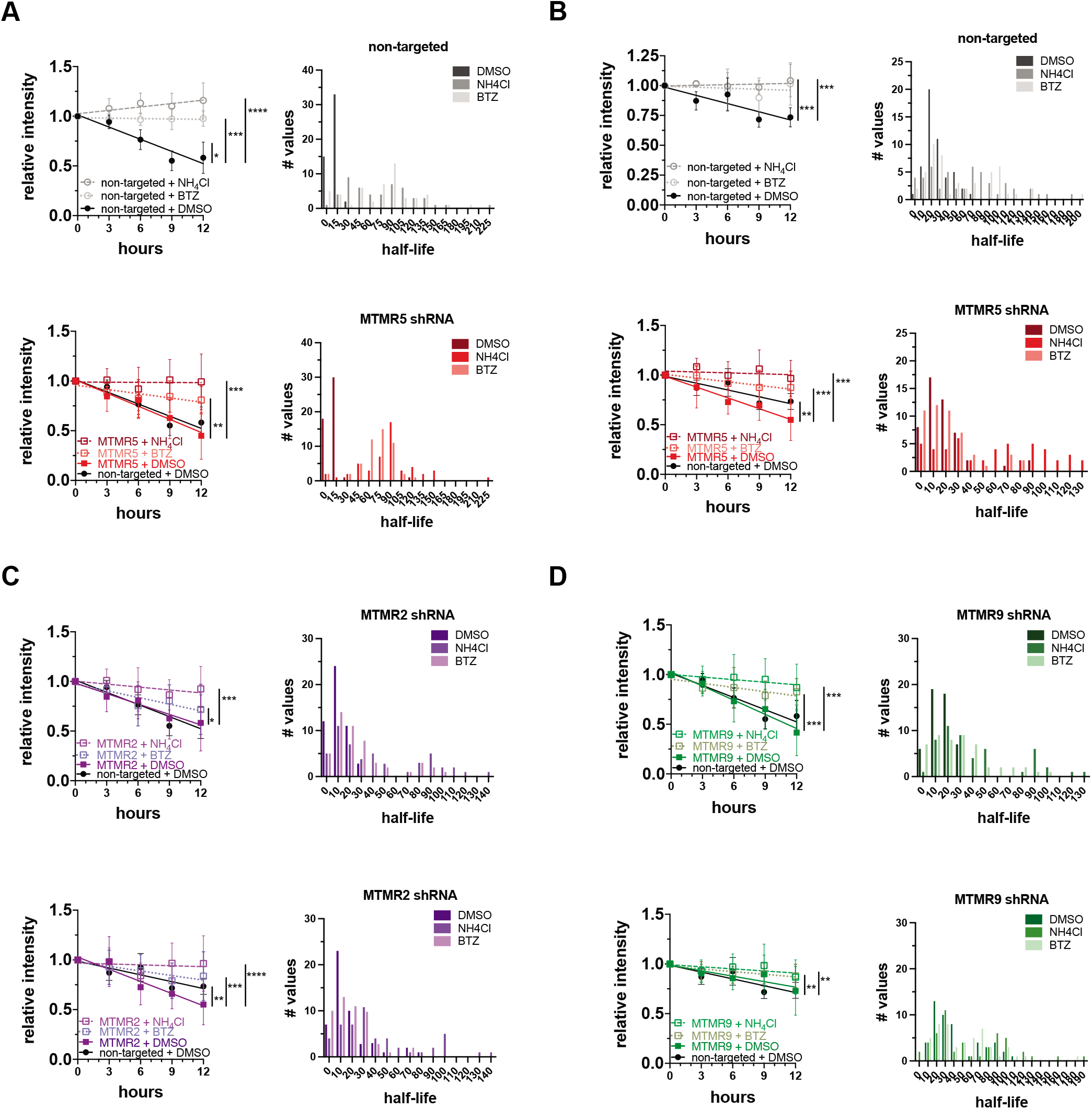
Degradation of autophagy substrates is not regulated by MTMR5-MTMR2 in iMotor Neurons. (**A**-**D**) Fluorescence of TDP-43-Dendra2 (A, C-D top) or Dendra2 (B, C-D bottom) measured in iMotor Neurons transduced with lentivirus expressing non-targeted shRNA (A-B, top panels) *SBF1* shRNA (A-B, bottom panels), *MTMR2* shRNA (C), or *MTMR9* shRNA (D) and after the indicated drug treatments (left panels), and histogram plot of the half-lives of each measured iMotor Neuron (right panels). Data are represented at each time point as mean ± SD; *p<0.05; **p<0.01; ***p<0.001; ***p<0.001; ****p<0.0001, one-way ANCOVA.

We also created GADPH-2A-Dendra2 iMotor Neurons to examine the turnover of Dendra2 alone by OPL in these cells. As with iNeurons, Dendra2 showed a half-life of about 24h in iMotor Neurons, and this was extended to similar degrees by ammonium chloride and bortezomib (Fig. 7B). *SBF1* and *MTMR2* knockdown both accelerated Dendra2 turnover in iMotor Neurons (Fig. 7B, 7C lower panels) while *MTMR9* knockdown had little effect (Fig. 7D, lower panel), testifying to the specificity of MTMR2/5-regulated autophagic function. These findings demonstrate not only proteolytic profiles unique to each individual degradative substrate of autophagy and specific for each neuronal subtype (Fig. S4G), but also cell type- and substrate-specific enhancement of autophagy after knocking down *SBF1* or *MTMR2*.

## DISCUSSION

In the present study, we identify a neuron-specific determinant of autophagy inhibition, MTMR5, the suppressive effects of which are dependent on MTMR2 (Fig. 6–7). Not only is *SBF1* mRNA stabilized in iNeurons compared to isogenic fibroblasts and iPSCs, but MTMR5 protein is also elevated in iNeurons. Overexpression of MTMR5 recapitulates neuron-like resistance to autophagy inducers in iPSCs, while reduction of MTMR5 or MTMR2 leads to enhanced autophagosome biogenesis, as well as accelerated clearance of two autophagy substrates, Dendra2-tagged TDP-43 and Dendra2 itself. These results highlight the MTMR5-MTMR2 axis as a novel means of selectively augmenting neuronal autophagy and enhancing the efficacy of autophagy-based therapies for neurodegenerative diseases (Fig. S5).

Our results also illustrate different profiles of autophagic activity in distinct cell types, correlating with each cell’s unique complement of autophagy-related factors. For instance, undifferentiated iPSCs are exquisitely sensitive to Torin1’s autophagy-inducing effects (Fig. 1E-G), and these cells exhibit low levels of *SBF1* (Fig. 2), but also higher levels of the autophagy-promoting factors *TFEB* (a master transcriptional regulator of autophagy and lysosomal genes) and *ATG5* (a critical component of the ATG5-ATG12-ATG16L1 complex which regulates phagophore elongation^88,89^) as determined by Bru-seq and RT-PCR (Fig. 2B-C, Fig. S6A). Of these, only *ATG5* knockdown (Fig. S6B) successfully desensitizes iPSCs to Torin1 and reduces autophagosome biogenesis (Fig. S6C-D). Interestingly, and to our surprise, iAstrocytes robustly generate autophagosomes after Torin1 treatment (Fig. 1E-G) despite expressing higher *SBF1* levels compared to their iPSCs of origin (Fig. S2C). One possible explanation for this observation is that iAstrocytes express lower levels of *MTMR2* (Fig. S2D) but also higher levels of *TFEB*, *ATG5*, and *SQSTM1* (Fig. S2E), the latter of which encodes the autophagy adaptor p62/sequestosome-1 that recruits ubiquitinated substrates to autophagosomes^29,90^. The combination of multiple autophagy-promoting factors may counteract the inhibitory influence *SBF1* in iAstrocytes, though the necessity of these factors for preventing neuron-like resistance to Torin1 remains to be determined.

Although iMuscle express similar levels of *SBF1* to iNeurons (Fig. S2B), Torin1 treatment potently induces autophagosome biogenesis in iMuscle (Fig. 1E-G). This discrepancy may relate to levels of *MTMR2* (Fig. S2C) or undetectable MTMR5 protein on immunocytochemistry staining (Fig. S1B), implying the presence of additional factors regulating MTMR5 translation in iMuscle. In line with this, pharmacologically or genetically abrogating autophagy leads to numerous skeletal muscle pathologies. After autophagy inhibition with 3-methyladenine, skeletal muscle in mice fails to recover and regrow following cardiotoxin injections^91^. Blocking autophagy through global knock-out of *ulk1*, an essential component of autophagy initiation machinery, constrains skeletal muscle regrowth after myectomy in zebrafish^92^. Conditional knock-out of *Atg5*^93^ or *Atg7*^94^ in mouse skeletal muscle leads to disruption of ultrastructural organization, accumulation of ubiquitinated inclusions, and dysfunctional mitochondria and sarcoplasmic reticulum. Thus, the abundance of pro-autophagy factors we noted in iMuscle may be necessary for promoting intact and productive autophagy, itself required for maintaining skeletal muscle proteostasis, organelle function, and structural integrity.

Separate from glutamatergic neurons, glia, and skeletal muscle, the regulatory mechanisms intrinsic to motor neurons deserve special mention. Not only are hallmarks of impaired autophagy such as ubiquitinated and p62-positive inclusions^16,95^ apparent in motor neuron disease, but also genetically attenuating *SBF1* and *MTMR2* has little effect in iMotor Neurons as compared to glutamatergic iNeurons. In addition, iMotor Neurons harbor high levels of *SBF1* (Fig. S4C) yet induce autophagosome formation to a greater extent than iNeurons upon Torin1 treatment (Fig. S4D-E). Notably, and as a potential counter to the suppressive influence of *SBF1*, iMotor Neurons express the highest levels of *TFEB* among the cell types analyzed, as well as higher levels of *ATG5* compared to iNeurons (Fig. S4F). The enhancing effects of these autophagy-promoting factors are reflected by shorter half-lives for TDP-43-Dendra2 and Dendra2, neither of which were further accelerated by *SBF1* or *MTMR2* knockdown (Fig. 7). Other components of the autophagy machinery in motor neurons have a profound influence on proteostasis and neuroprotection, as demonstrated by *Atg7*^96^ and *Tbk1*^97^ conditional deletion in motor neurons leading to earlier neurodegeneration in transgenic mouse models of amyotrophic lateral sclerosis. Interestingly, attenuated autophagy in these studies limited glial inflammation and extended lifespan^96,97^, emphasizing the interplay of cell and non-cell autonomous effects of disrupting autophagy on disease pathogenesis. These effects may also be species-specific, given that murine embryonic stem cell (ESC)-derived cranial motor neurons harbor greater proteasomal capacity and resistance to protein aggregation-related toxicity in comparison to ESC-derived spinal motor neurons^98^. Regardless, our results not only suggest that human iMotor Neurons operate at a higher basal rate of autophagic flux, but also that individual neuronal subtypes may be unique in their individual complement^99^ of activating factors and, in turn, intrinsic rates of autophagic activity. Such differences may explain discrepant reports in the literature regarding the extent to which neurons respond to autophagy stimuli^31,37,52,54–56,74^, not just in cell and tissue samples containing multiple subtypes of neurons, but also those containing heterogenous populations of non-neuronal cells.

One function of myotubularins is to modulate steady-state pools of phosphoinositides by reducing PtdIns3P and PtdIns(3,5)P2^59,100^. Our results imply that targeting MTMR5 is a promising strategy for amplifying autophagic degradation of misfolded proteins in neurodegenerative disorders, but additional aspects of MTMR5 function require further clarification. First, the relative levels of phosphoinositides and their subcellular distribution in neurons^52,101–103^ after *SBF1* or *MTMR2* knockdown may confirm the necessity of MTMR5 potentiating the phosphatase activity of MTMR2, especially at neuronal phagophore assembly sites. Second, given the presence of additional protein-protein interaction motifs in MTMR5, including PH (plekstrin-homology), DENN (differentially expressed in normal and neoplastic cells), and SID (SET-interacting domain) motifs, and other documented MTMR5 binding partners including multiple Rab GTPases^104,105^ and *KMT2A*/HRX^106^, it remains a possibility that MTMR5 modulates autophagy through additional or alternative mechanisms such as membrane trafficking^104,105^ or transcriptional and epigenetic regulation^107,108^. Third, it remains unknown if targeting MTMR5 is the optimal strategy for rescuing protein dyshomeostasis in the central nervous system while avoiding peripheral and age-dependent requirements of intact MTMR5. Ideally, newly developed therapies would not reduce peripheral MTMR5 expression, since loss of testicular *Sbf1* in mice leads to infertility^109^, and dysfunction in Schwann cell MTMR5 leads to CMT4B3^64^, a dysmyelinating inherited polyneuropathy. Fourth, the mechanisms by which the *SBF1* transcript is stabilized in iNeurons remain undefined. Possibilities include hypomethylation of neuronal *SBF1*^110^ or neuronal enrichment of enhancers with cognate binding sites in the *SBF1* promoter region^111,112^, including BRD3, NOTCH1, and TCF12^113^. However, our Bru-Seq data show that synthesis of *SBF1* transcripts is similar between iNeurons and iPSCs (Fig. 2B), suggesting that heightened *SBF1* stability (Fig. 2B-E) may instead be due to neuron-specific attenuation of *SBF1* decay. Lastly, what are the teleologic underpinnings of MTMR5 expression and restricted autophagy in neurons? It is possible that limiting excess autophagy induction in neurons affords an evolutionary advantage by preventing depletion of essential cytoplasmic constituents^114^ or exacerbating inefficient cargo capture that outstrips the degradative capacity of the lysosomal compartment in neurons^53,115,116^, though these possibilities require further investigation. Nonetheless, our results establish a critical role of MTMR5 in neuronal autophagy and identify a novel and promising mechanistic target for ameliorating proteostatic deficits in neurodegenerative diseases.

## ACKNOWLEDGEMENTS

We thank Sandra Mojica-Perez and Michael Uhler (University of Michigan Human Stem Cell and Gene Editing Core) and Andrea Serio (The Francis Crick Institute) for their expertise and guidance in iAstrocyte differentiation. Figures 1A-D, 2A, 3A, 5A-B, 5F, and S5 were created using BioRender (https://www.biorender.com). This research was supported by the NIH National Institute for Neurological Disorders and Stroke (NINDS) R25 NS089450-06, R01 NS099340, and the Intramural Research Program of the NIH/NINDS.

## AUTHOR CONTRIBUTIONS

Conceptualization, J.P.C., L.S.W., and S.J.B.; Methodology, J.P.C., K.B., M.T.P., M.L., E.M.H.T., E.S.K., J.M.C.M., M.E.W., L.S.W., and S.J.B.; Software, S.J.B.; Formal Analysis, J.P.C., K.B., M.T.P., M.L., and S.J.B.; Investigation, J.P.C., E.M.H.T., E.S.K., and S.J.B.; Resources, J.M.C.M. and M.E.W.; Writing – Original Draft, J.P.C. and S.J.B.; Writing – Review & Editing, J.P.C., K.B., M.T.P., M.L., E.M.H.T., E.S.K., J.M.C.M., M.E.W., L.S.W., and S.J.B.; Visualization, J.P.C., K.B., M.T.P., M.L., L.S.W., and S.J.B.; Supervision, L.S.W.; Project Administration, J.P.C., E.M.H.T., and S.J.B.; Funding Acquisition, J.P.C., J.M.C.M., M.E.W., L.S.W., and S.J.B.

## MATERIALS AND METHODS

### Generation of iPSC lines

Human iPSCs reprogrammed from skin fibroblasts obtained from a healthy adult male were engineered by the Allen Institute for Cell Science (https://www.allencell.org) to express fluorescently labeled LC3 using CRISPR/Cas9 to insert EGFP at the N-terminal exon of the *MAP1LC3B* gene. These iPSCs were then engineered for rapid differentiation into neuronal cell types using TALENS to introduce doxycycline-inducible transcription factors at the *CLYBL* safe harbor locus, either neurogenin-1 and −2 (*NGN1/2*) for generating forebrain-like cortical neurons (iNeurons), or *LHX3/ISL1/NGN2* for generating of motor neurons (iMotor Neurons). For rapid differentiation into iMuscle, cells were genomically edited with piggy-bac/transposon vectors kindly provided by Michael Ward (NINDS) and containing doxycycline-inducible MYOD1/shRNA-OCT4.

### Differentiation protocols

Rapid differentiation was performed as previously described^6^. Briefly, on DIV 0 iPSCs were split with Accutase and plated on PEI-coated slides for live cell imaging or Matrigel-coated plates for RNA and protein analyses. For iNeurons, on DIV 1-2 media were changed daily from N2 medium (TesR-E8) to transition medium (half TesR-E8, half DMEM/F12), with both media supplemented with doxycycline, N2, NEAA, BDNF, NT3, and laminin. On DIV 3, media were changed to B27 medium (Neurobasal-A supplemented with doxycycline, B27, Glutamax, BDNF, NT3, laminin, and Culture One). From DIV 6-14, equal volumes of B27 medium without Culture One were added every 4 days to each well. For iMotor Neurons, media were changed on DIV 1 to D1 medium (TesR-E8 supplemented with doxycycline, N2, and Compound E) and DIV 3 to D3 medium (DMEM/F12 supplemented with doxycycline, N2, NEAA, Glutamax, and Compound E). On DIV 6, media were changed to B27 medium with Culture One. From DIV 10-14, equal volumes of B27 medium without Culture One were added every 4 days to each well. For iAstrocytes^68^, iPSCs were differentiated to neuroectoderm by dual-SMAD signaling inhibition in 3N medium (half Neurobasal-A, half DMEM/F12 supplemented with SB431542, dorsomorphin, N2, B27, NEAA, Glutamax, hr-insulin, and pen/strep) for 2-4 weeks. Following this, neurospheres were formed and mechanically chopped, passaged, and cultured in EL20 medium (half Neurobasal-A, half DMEM/F12 supplemented with EGF, LIF, N2, B27, NEAA, Glutamax, and pen/strep) for 2-4 weeks. Cells were then passaged and maintained as monolayers of astrocyte precursors in EF20 medium (half Neurobasal-A, half DMEM/F12 supplemented with EGF, FGF2, N2, B27, NEAA, and Glutamax) for 2-4 weeks. Finally, astrocyte precursors were differentiated to mature astrocytes with AstroMED medium (Neurobasal-A supplemented with CNTF, B27, NEAA, Glutamax, and pen/strep) for 4 weeks. For iMuscle, iPSCs were cultured in myogenic progenitor media (MEMa supplemented with sodium pyruvate, non-essential amino acids, Glutamax, 2-mercaptoethanol, doxycycline, and 5% knock-out serum replacement) for 2 days, followed by myogenic induction medium (DMEM supplemented with Glutamax, IGF-1, 2-mercaptoethanol, non-essential amino acids, doxycycline, and 5% knock-out serum replacement) for an additional 5 days.

### Primary cell culture

Rat cortical neurons and cortical glia were dissected from E20-E21 rat pups, fractionated by centrifugation, and plated on poly-D-lysine (PDL)-coated flasks at a density of ~30-50 × 10^6^ cells per flask. Cells were maintained with media changes every 2 days. After DIV 7-8, glial flasks were shaken with a benchtop cell agitator at 180rpm x 30min to detach and remove microglia, then 240rpm for 6 hours and washed twice with PBS to detach and remove oligodendrocyte precursors. Purified cortical astrocytes were then dissociated with 0.25% Trypsin-EDTA, re-plated in new PDL-coated flasks, maintained in culture for 12-14 days, then collected for RNA and protein analyses.

### Study approval and ethics statement

All vertebrate animal work was approved by the Committee on the Use and Care of Animals (UCUCA) at the University of Michigan and in accordance with the United Kingdom Animals Act (1986). All experiments were performed in accordance with UCUCA guidelines. Rats (Rattus norvegicus) used for primary neuron collection were housed singly in chambers equipped with environmental enrichment. All studies were designed to minimize animal use. Rats were cared for by the Unit for Laboratory Animal Medicine at the University of Michigan; all individuals were trained and approved in the care and long-term maintenance of rodent colonies, in accordance with the NIH-supported Guide for the Care and Use of Laboratory Animals (National Academies Press, 2011). All personnel handling the rats and administering euthanasia were properly trained in accordance with the University of Michigan Policy for Education and Training of Animal Care and Use Personnel. Euthanasia was performed according to the recommendations of the Guidelines on Euthanasia of the American Veterinary Medical Association.

### Transfection

Prior to transfection, iPSCs were split with EDTA and seeded onto vitronectin-coated plates. On the day of transfection, cells were switched from TeSR E8 (StemCell Technologies 05990) to mTeSR 1 (StemCell Technologies 85850), and total DNA amounts of 5μg (2.5μg donor vector plus 1.25μg of each TALENS arm plasmid) or 2μg (1μg piggybac vector plus 1μg of transposase plasmid) were combined with Lipofectamine Stem (ThermoFisher, STEM00003) in Opti-MEM (ThermoFisher 11058021), then added dropwise to iPSCs and incubated overnight, followed by media change to TeSR-E8 the next morning.

### Lentiviral transduction

TRC Lentiviral shRNA plasmids against human *SBF1* (Horizon RHS3979-201739167), *MTMR2* (Horizon RHS3979-224867040), *MTMR9* (Horizon RHS3979-201908246), *TFEB* (Horizon RHS3979-201744686), *ATG5* (Horizon RHS3979-201857832), *SQSTM1* (Horizon RHS3979-201739507), or non-targeted control (Horizon RHS6848) were produced through the University of Michigan Vector Core. On DIV7, half the working volume of B27 medium was removed from iNeuron and iMotor Neuron cultures and saved at 4º C, or TeSR E8 from undifferentiated iPSCs, and replaced with equivalent volumes of lentiviral lysate. The following day, the saved conditioned B27 media was re-warmed at room temperature and added back to iNeuron and iMotor Neuron cultures. The remaining seven days of differentiation were continued unchanged and as described above, and cells were imaged on DIV14. For iPSCs, fresh TeSR-E8 was added the following day after addition of lentiviral lysates, and daily media changes continued for 6 more days before treatment with Torin1 and imaging.

### Bru-seq and BruChase-seq

RNA labeled with bromouridine (BrU) was prepared and analyzed as previously described^117^. Briefly, BrU-labeled RNA was extracted with anti-BrU antibodies from total RNA samples from healthy adult fibroblasts, isogenic iPSCs reprogrammed from these fibroblasts, and isogenic iNeurons differentiated from these iPSCs. Strand-specific DNA libraries were prepared with Illumina TruSeq Kit (Illumina) and sequenced using the Illumina sequencing platform24. Strand-specific sequenced data was aligned to human ribosomal DNA complete repeating unit (U13369.1) using Bowtie^118^ (v0.12.8). Remaining unaligned reads were then mapped to the human genome build hg19/GRCh37 using TopHat^119^ (v1.4.1)24. Bru-seq data from iPSCs and iNeurons were compared to fibroblasts and fold differences quantified using DESeq^120^ (version 1.4.1) in R^121^ (version 2.15.1). Genes having a mean RPKM ≥ 0.5, length ≥ 300 bp, false discovery rate (FDR) ≤ 0.1 and a 1.5-fold change were included for downstream bioinformatics analyses. For BruChase-seq, a stability index for each transcript was calculated as a ratio of transcript abundance at 6 hours vs. 0.5 hours, and median values across replicates of the same condition were used. Genes showing greater or less than 1.5-fold change in the stability index were included in final dataset.

### Live cell imaging and optical pulse labeling

Cells were incubated in maintenance medium for each respective cell type supplemented with DMSO vehicle or 250nM Torin1 for four hours. For autophagosome puncta quantification, live cells were imaged using ONI Nanoimager (https://www.oni.bio), and images were processed and analyzed using Fiji. For optical pulse labeling, an automated microscopy platform was used as previously described^74,122,123^. Briefly, images were obtained at the indicated time points with a Nikon TE2000 microscope equipped with the PerfectFocus system, a high-numerical aperture 20× objective lens and a 16-bit Andor Clara digital camera with a cooled charge-coupled device. Illumination was provided by a Lambda XL Xenon lamp (Sutter) with a liquid light guide. The ASI 2000 stage was controlled by rotary encoders in all three planes of movement. All components were housed in a custom-designed, climate-controlled environmental chamber (InVivo Scientific) kept at 37° C and 5% CO_2_. The Semrock BrightLine full-multiband filter set (DAPI, FITC, TRITC, Cy5) was used for fluorophore photoactivation (DAPI), excitation and detection (FITC, TRITC). The illumination, filter wheels, focusing, stage movements and image acquisitions were fully automated and coordinated with a mix of proprietary (ImagePro) and publicly available (ImageJ^124^ and μManager^125^) software.

### RT-PCR

RNA was isolated from cell pellets using the RNeasy Mini Kit (QIAgen). cDNA was reverse transcribed from 1μg RNA with the iScript kit (Bio-Rad). Quantitative PCR was then performed on 0.5 μl of cDNA using Power SYBR Green (Applied Biosystems) with primers complementary to human *SBF1, MTMR2*, *TFEB*, *ATG5*, *SQSTM1*, and *GAPDH* for iPSCs and differentiated cells, or to rat *Sbf1* and *Gapdh* for primary cultured cells.

### Western blot

Cell pellets were lysed in ice-cold RIPA buffer supplemented with c*0*mplete protease inhibitors (Roche). Whole-cell lysates were then sonicated and clarified by centrifugation at 14,000xg for 15 minutes at 4°C. Protein concentrations were determined by BCA protein assay. After boiling at 100°C for 10 minutes in loading buffer, 20μg of protein samples were resolved on 10% or 15% SDS-PAGE gels, transferred to 0.2-μm PVDF membranes using the Mini Trans-Blot system (Bio-Rad), and probed by the indicated antibodies. Detection was performed by chemiluminescence.

### Immunocytochemistry

Cells were fixed in 4% paraformaldehyde (Sigma P6148) and permeabilized with 0.1% Triton X-100 (Bio-Rad 161-0407) in 1X phosphate buffered saline (PBS). Fixative was quenched with 10mM glycine (Fisher BP381-1), followed by incubating in blocking solution comprised of 3% bovine serum albumin w/v (BSA, Fisher BP9703-100), 2% fetal calf serum v/v (Sigma F4135), and 0.1% Triton X-100 v/v in 1X PBS for one hour at room temperature. Cells were then incubated with the indicated primary antibodies and diluted in blocking solution at 4º C overnight, washed three times for five minutes each in 1X PBS, then incubated with the indicated secondary antibodies (Table 2) diluted in blocking solution at room temperature for one hour. Finally, cells were washed three times for five minutes each in 1X PBS, washed twice for five minutes each in Hoescht 33258 dye (Invitrogen H3569, 1:10000 in 1X PBS), then mounted for fluorescence imaging.

### Quantification and statistical analysis

All live cell imaging, RT-PCR, and Western blot experiments were performed in biological triplicate and quantified using Fiji^126^. Quantification of mEGFP-LC3-positive puncta were blinded to genotype and drug treatment. Quantifications of image analyses and Western blot band intensities, normalized to actin loading control, are reported as mean ± SEM. Statistical significance of mean differences was determined using Student’s unpaired *t* test, or multiple comparisons with ANOVA and ANCOVA, and with p<0.05 considered significant. Data were analyzed and plotted using GraphPad Prism.

