## Supplemental materials for "Myotubularin-related phosphatase 5 is a critical determinant of autophagy in neurons"

**Figure S1. Generation of inducible myocytes from iPSCs.** (**A**) Immunocytochemical staining of muscle-specific markers Pax7, MF20, and MyoD in iMuscle and isogenic iPSCs. (**B**) Immunocytochemical staining for MTMR5 protein in iMuscle and their iPSCs of origin.

**Figure S2. Cell-type specific expression of autophagy-related effectors.** (**A**) Expression of *SBF1* in iNeurons at the indicated days of differentiation *in vitro* (iNeuron RNA-Seq app, Kampmann Laboratory[127]). (**B**) Expression of human *SBF1* in different neuronal subtypes and glia (Human – Multiple Cortical Areas Smart-seq Data Set, Allen Brain Atlas, https://celltypes.brain-map.org/rnaseq/human_ctx_smart-seq). (**C**) RT-PCR measurements of total steady-state *SBF1* RNA in iNeurons compared to iAstrocytes and iMuscle. n.s., not significant; ***$p<0.001$, one-way ANOVA. (**D**) RT-PCR measurements of human *MTMR2* or rat *Mtmr2* RNA in the indicated cell types. n.s., not significant; *$p<0.05$; **$p<0.01$, one-way ANOVA. (**E**) RT-PCR measurements of human *TFEB*, *ATG5*, or *SQSTM1* RNA in iNeurons compared to iAstrocytes; **$p<0.01$; ***$p<0.001$; ****$p<0.0001$, Student's *t* test.

**Figure S3. Knockdown of MTMR2 sensitizes neurons to Torin1.** (**A**) Representative super resolution images of iNeurons transduced with non-targeted shRNA or *MTMR2* shRNA and treated with DMSO vehicle or 250nM Torin1 for four hours. (**B**) Scatterplots of blinded manual quantifications of mEGFP-LC3-positive puncta imaged in iNeurons as treated in (A). Data are from three independent experiments; n.s., not significant, Student's *t* test. (**C**) Histogram plots of mEGFP-LC3-positive puncta quantifications from (B).

**Figure S4. iMotor Neurons have more robust autophagy induction than iNeurons.** (**A**) Schematic of the cassette used to integrate *LHX3*, *ISL1*, and *NGN2* at the *CLYBL* safe harbor locus under the control of a Tet-ON system. *Neo*, neomycin-resistance gene; *pA*, poly-A tail; $P_1$, $P_2$, promotors; *iRFP*, near-infrared fluorescent protein; *rTTA*, reverse tetracycline-controlled transactivator; *NGN1* and *NGN2*, neurogenin-1 and -2; *T2A*, self-cleaving peptide; *TRE*, tetracycline response element. (**B**) Immunocytochemical staining of DIV14 iMotor Neurons for motor neuron markers P75, choline acetyltransferase (ChAT), and MAP2. (**C**) RT-PCR measurements of human *SBF1* RNA in iMotor Neurons compared to isogenic iPSCs and iNeurons; n.s., not significant; **p<0.01; ***p<0.001, one-way ANOVA. (**D**) Representative images of mEGFP-LC3-positive vesicles in iNeurons and iMotor Neurons after treatment with DMSO vehicle or 250nM Torin1 for 4 hours. (**E**) Scatterplots of blinded manual quantifications of mEGFP-LC3-positive vesicles imaged as in (D). Data are from three independent experiments; n.s., not significant; *p<0.05; **p<0.01, Student's *t* test. (**F**) RT-PCR measurements of human *TFEB* and *ATG5* RNA in iMotor Neurons compared to iNeurons; *p<0.05; ****p<0.0001, Student's *t* test. (**G**) Dendra2 fluorescence measured in TDP-43-Dendra2 (left) and GAPDH-2A-Dendra2 (right) iNeurons and iMotor Neurons transduced with non-targeted shRNA lentivirus, to compare endogenous rates of decay for the substrates TDP-43 and Dendra2 in each cell type; n.s., not significant ****p<0.0001, one-way ANCOVA.

**Figure S5. Proposed model of neuron-specific regulation of autophagy.** (**A**) In non-neuronal cells (left), autophagy operates under permissive conditions due to relatively lower levels of MTMR5, enabling sufficient levels of PtdIns3P and PtdIns(3,5)P2 to recruit autophagy-related protein complexes (ATG machinery) in response to stimuli (e.g., Torin1) and necessary for autophagosome biogenesis. However, in neurons (right), higher levels of MTMR5 impair autophagy induction by potentiating MTMR2 phosphatase activity to deplete the phosphoinositides needed for ATG machinery recruitment. (**B**) In neurons, autophagy induction is suppressed by native levels of MTMR5 (left), but opposing MTMR5 activity (e.g., through shRNA-mediated knockdown) disinhibits autophagy and accelerates autophagic degradation of substrates such as TDP-43 (right).

**Figure S6. ATG5, but not TFEB, is necessary for maintaining sensitivity to Torin1.** (**A**) RT-PCR measurements of human *TFEB* and *ATG5* RNA in iNeurons compared to isogenic iPSCs; *p<0.05; ***p<0.001, Student's *t* test. (**B**) Representative immunocytochemical staining against TFEB (left) or ATG5 (right) in iPSCs after transduction with non-targeted, *TFEB*, or *ATG5* shRNA lentivirus. Scale bar, 30μm. (**C**) Representative images of mEGFP-LC3-positive vesicles in iPSCs after treatment with DMSO vehicle or 250nM Torin1 for 4 hours and transduction with non-targeted (NT), *TFEB*, or *ATG5* shRNA lentivirus (left), and scatterplots (right) of blinded manual quantifications of mEGFP-LC3-positive vesicles. Scale bar, 5μm. Data are from three independent experiments; n.s., not significant; ****p<0.0001; one-way ANOVA with Šídák's multiple comparisons test.

Figure S1

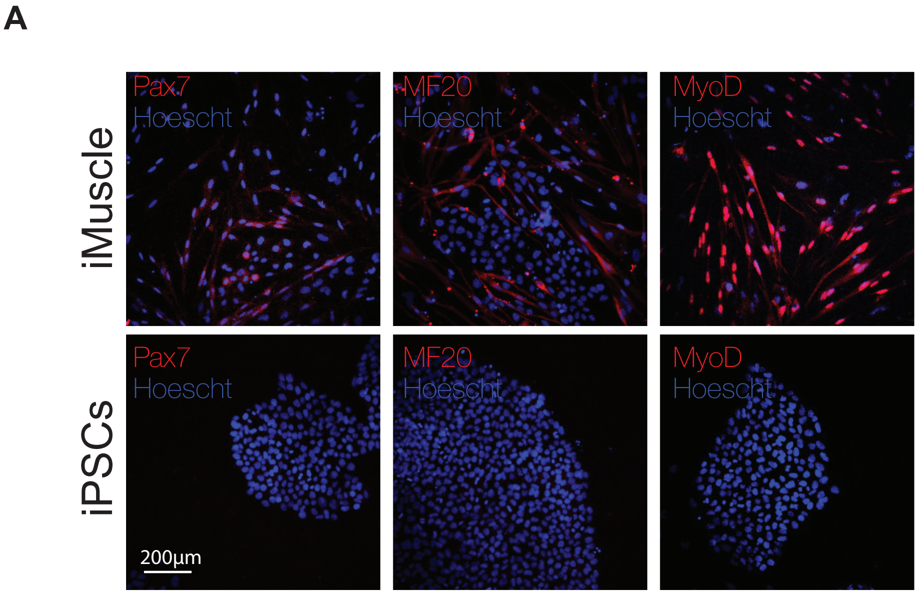
A
iMuscle
iPSCs
Pax7
Hoescht
MF20
Hoescht
MyoD
Hoescht
Pax7
Hoescht
MF20
Hoescht
MyoD
Hoescht
200μm

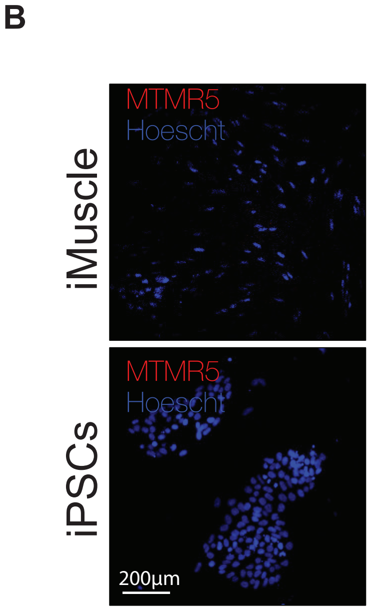
B
iMuscle
iPSCs
MTMR5
Hoescht
MTMR5
Hoescht
200μm

### Figure S2

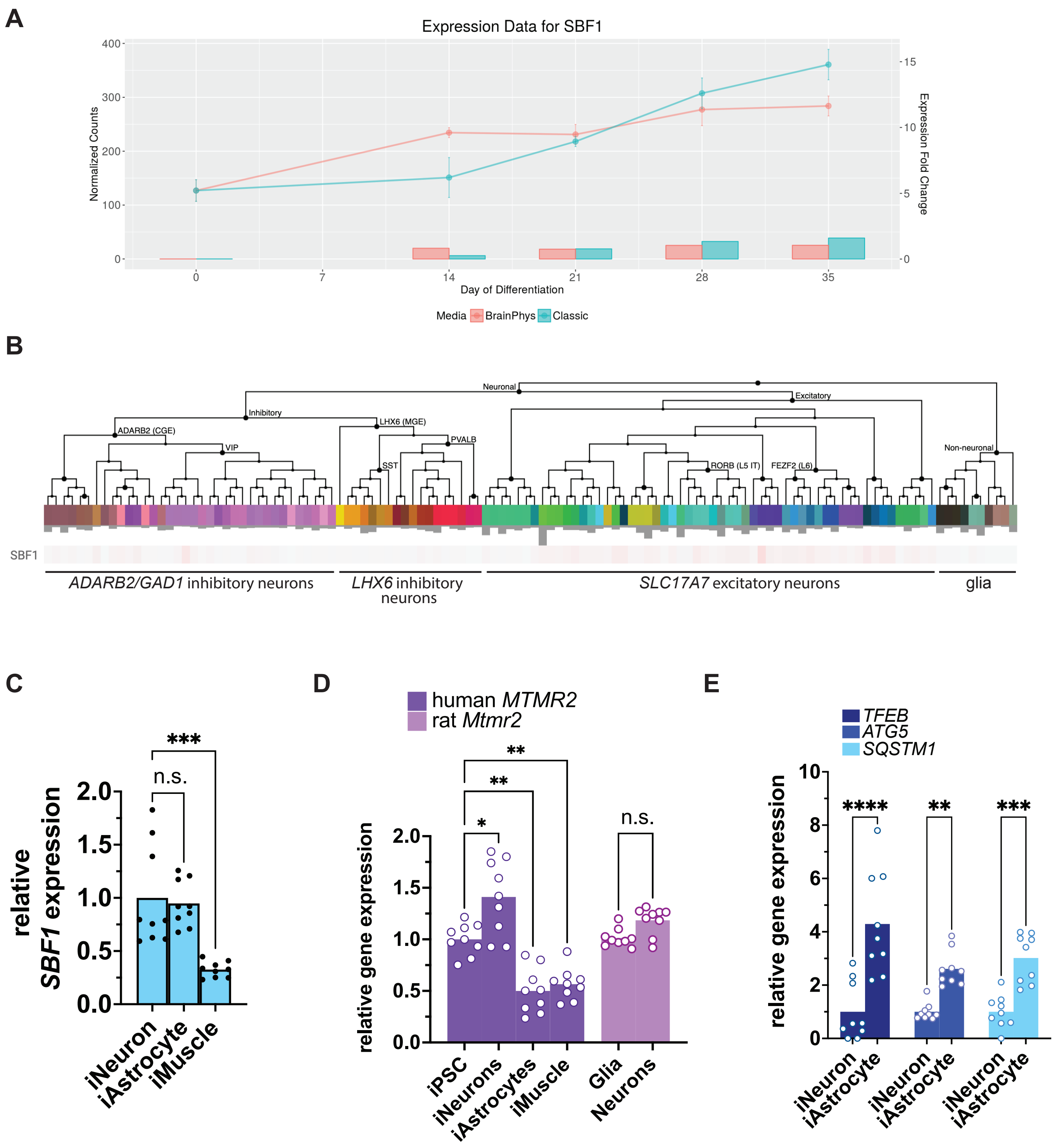
A
Expression Data for SBF1
Normalized Counts
Expression Fold Change
Day of Differentiation
Media
BrainPhys
Classic
B
Neuronal
Inhibitory
Excitatory
ADARB2 (CGE)
LHX6 (MGE)
VIP
PVALB
SST
RORB (L5 IT)
FEZF2 (L6)
Non-neuronal
SBF1
ADARB2/GAD1 inhibitory neurons
LHX6 inhibitory neurons
SLC17A7 excitatory neurons
glia
C
relative SBF1 expression
n.s.
***
iNeuron
iAstrocyte
iMuscle
D
human MTMR2
rat Mtmr2
relative gene expression
*
**
**
n.s.
iPSC
iNeurons
iAstrocytes
iMuscle
Glia
Neurons
E
TFEB
ATG5
SQSTM1
relative gene expression
****
**
***
iNeuron
iAstrocyte

Figure S3

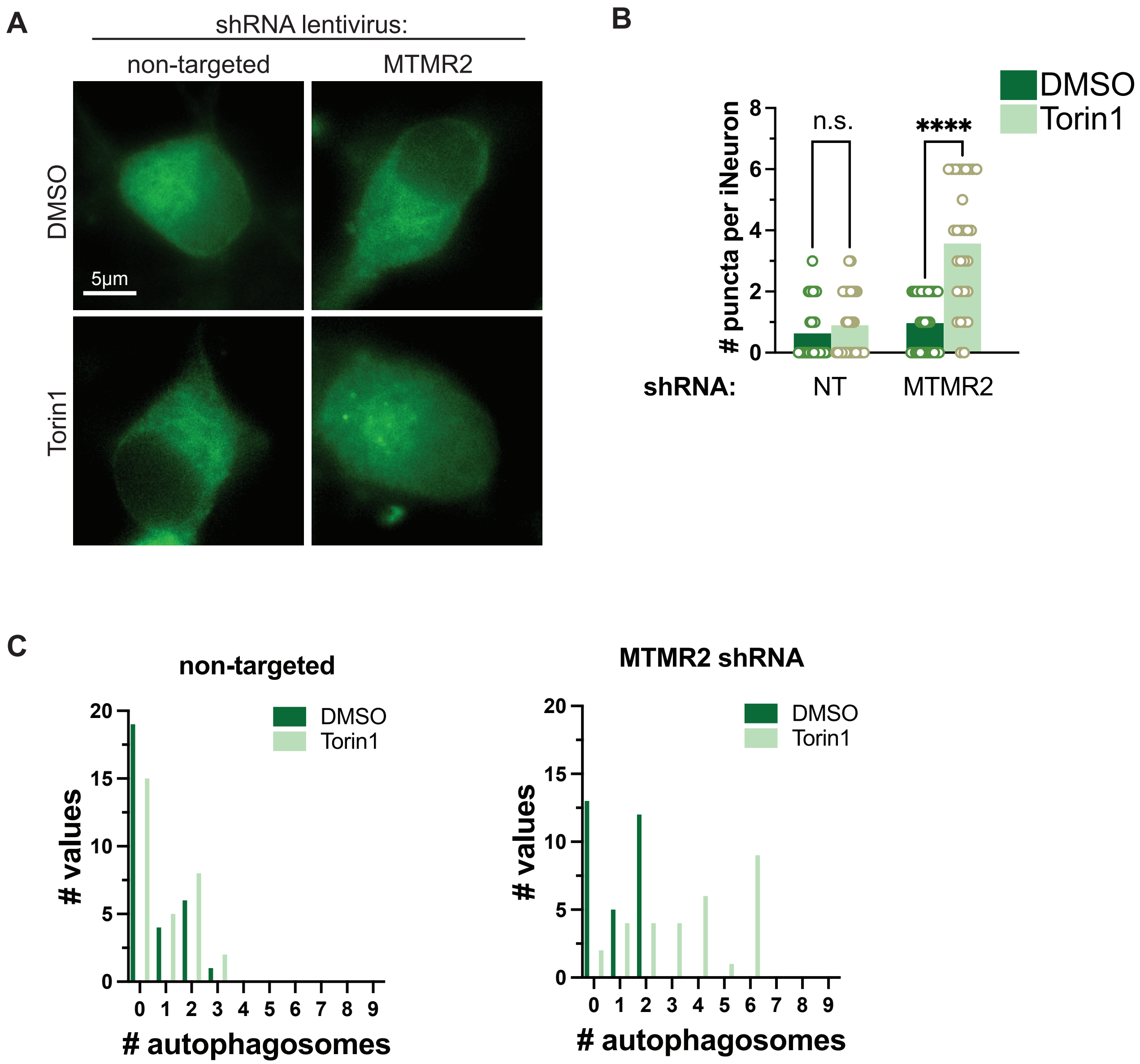
A
shRNA lentivirus:
non-targeted
MTMR2
DMSO
Torin1
5µm
B
DMSO
Torin1
### puncta per iNeuron
n.s.
****
shRNA:
NT
MTMR2
C
non-targeted
MTMR2 shRNA
### values
### autophagosomes
DMSO
Torin1

### Figure S4

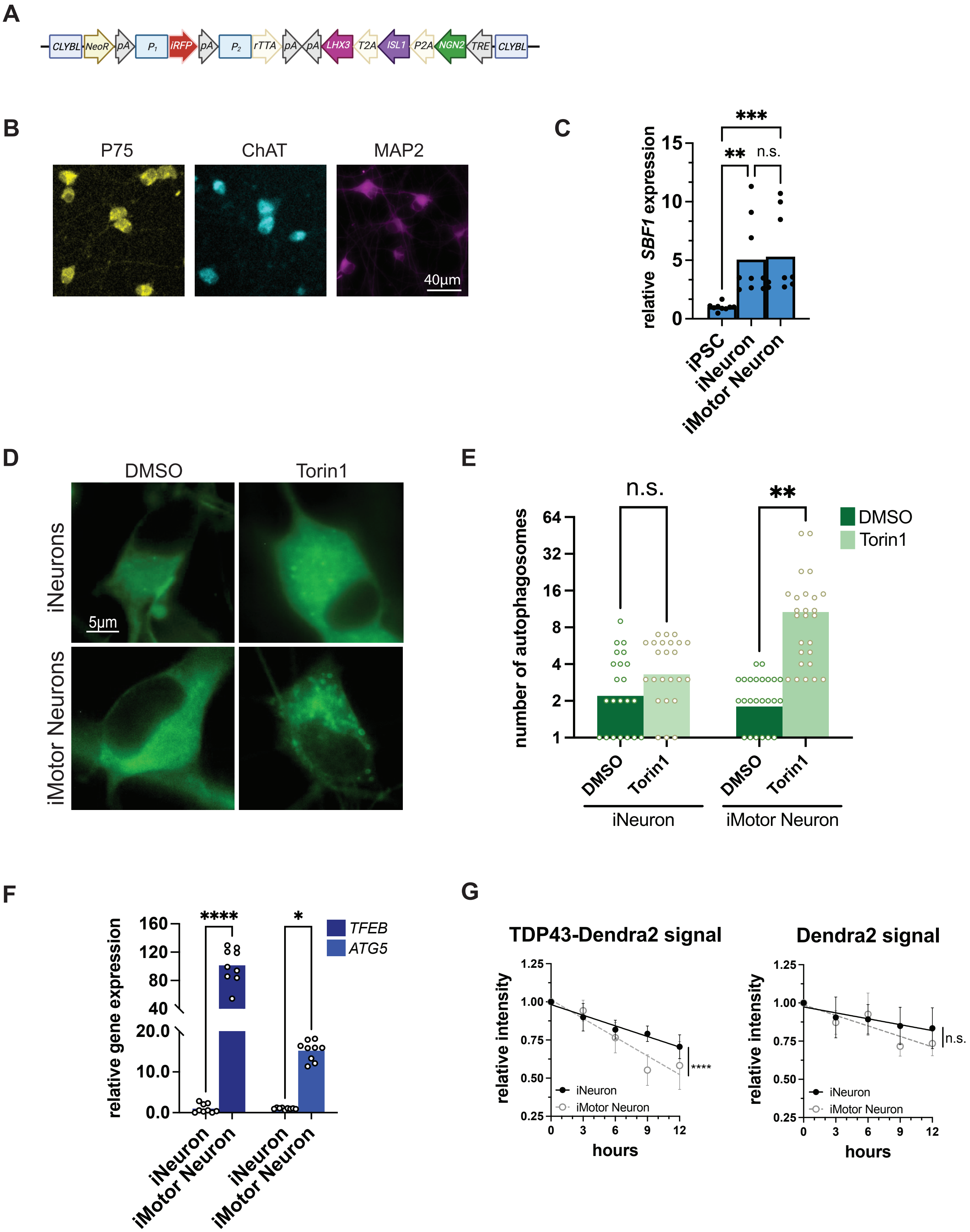

A
CLYBL
NeoR
pA
P1
iRFP
pA
P2
rTTA
pA
pA
LHX3
T2A
ISL1
P2A
NGN2
TRE
CLYBL
B
P75
ChAT
MAP2
40μm
C
relative SBF1 expression
***
**
n.s.
iPSC
iNeuron
iMotor Neuron
D
DMSO
Torin1
iNeurons
iMotor Neurons
5μm
E
number of autophagosomes
n.s.
**
DMSO
Torin1
iNeuron
iMotor Neuron
F
relative gene expression
****
*
TFEB
ATG5
iNeuron
iMotor Neuron
iNeuron
iMotor Neuron
G
TDP43-Dendra2 signal
Dendra2 signal
relative intensity
hours
iNeuron
iMotor Neuron
****
n.s.

### Figure S5

**A**

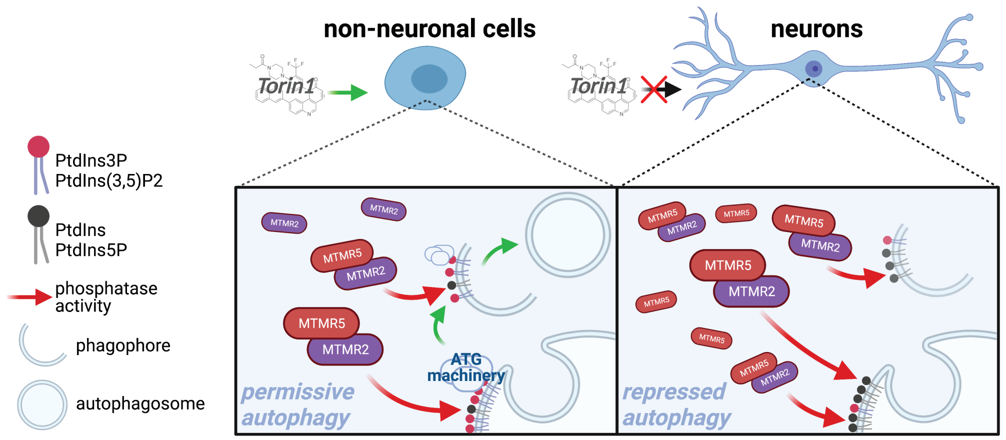
non-neuronal cells
neurons
Torin1
Torin1
PtdIns3P
PtdIns(3,5)P2
PtdIns
PtdIns5P
phosphatase activity
phagophore
autophagosome
MTMR2
MTMR5
ATG machinery
permissive autophagy
repressed autophagy

**B**

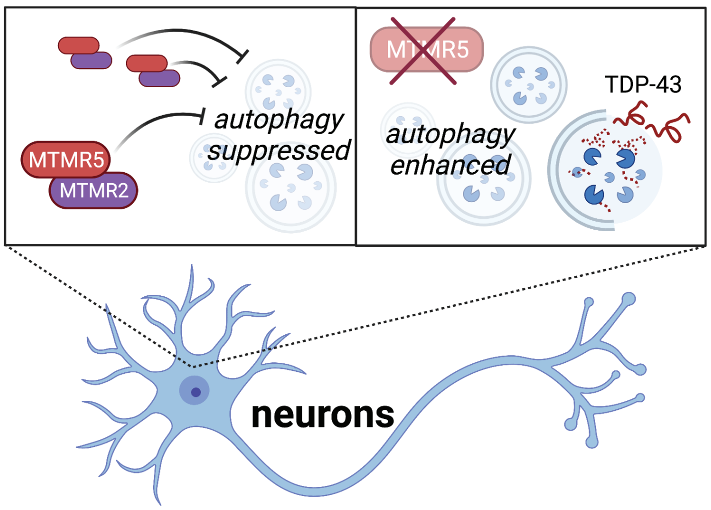
MTMR5
MTMR2
autophagy suppressed
MTMR5
autophagy enhanced
TDP-43
neurons

### Figure S6

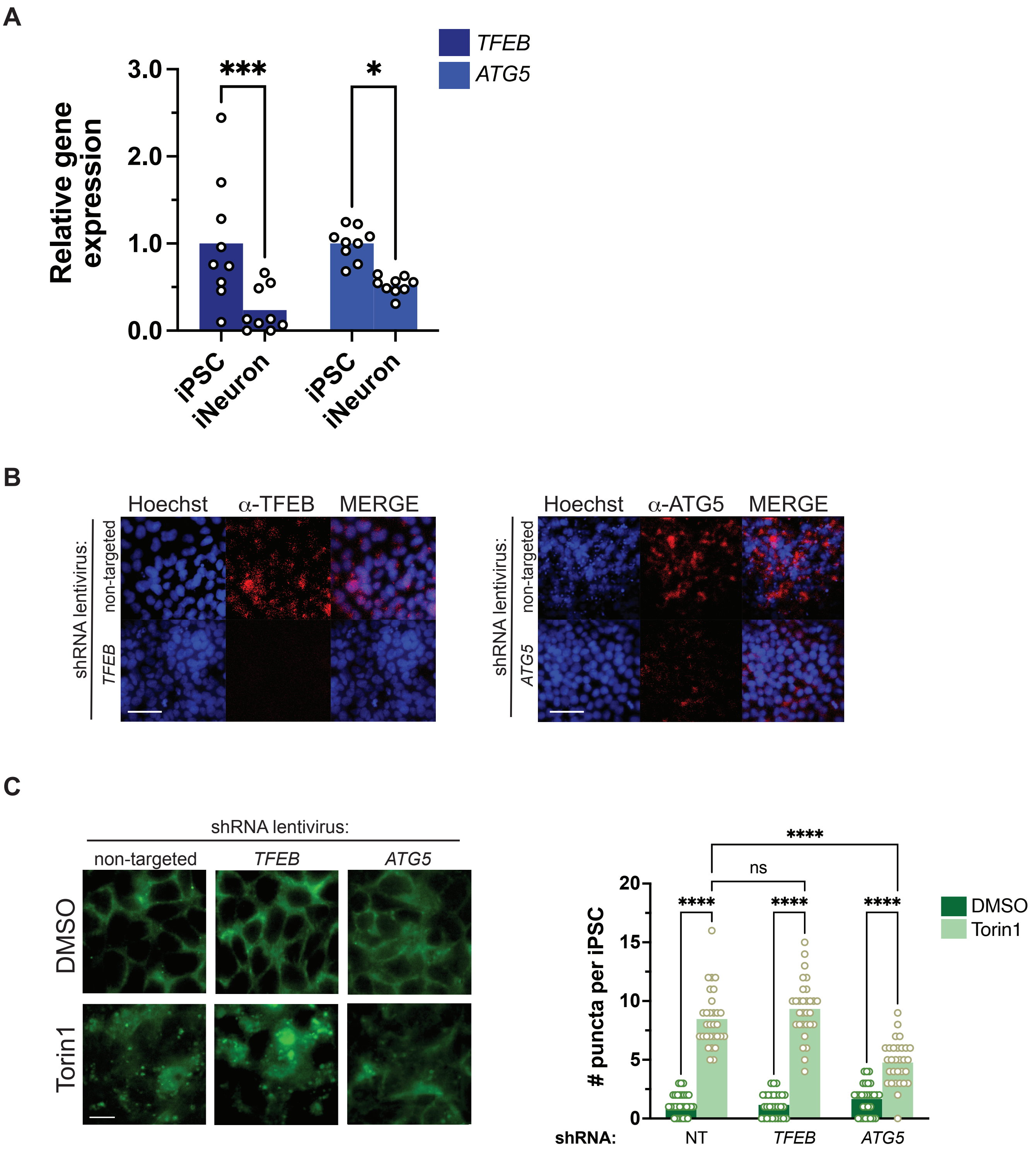

A
TFEB
ATG5
Relative gene expression
3.0
2.0
1.0
0.0
***
*
iPSC
iNeuron
iPSC
iNeuron
B
Hoechst
α-TFEB
MERGE
shRNA lentivirus:
non-targeted
TFEB
Hoechst
α-ATG5
MERGE
shRNA lentivirus:
non-targeted
ATG5
C
shRNA lentivirus:
non-targeted
TFEB
ATG5
DMSO
Torin1
### puncta per iPSC
20
15
10
5
0
****
ns
****
****
****
DMSO
Torin1
shRNA:
NT
TFEB
ATG5
